# Strong pairwise interactions do not drive interactions in a plant leaf associated microbial community

**DOI:** 10.1101/2024.05.22.595276

**Authors:** Franziska Höhn, Vasvi Chaudhry, Caner Bagci, Maryam Mahmoudi, Elke Klenk, Lara Berg, Paolo Stincone, Chambers C. Hughes, Daniel Petras, Heike Brötz-Oesterhelt, Eric Kemen, Nadine Ziemert

**Affiliations:** Translational Genome Mining for Natural Products, Interfaculty Institute of Microbiology and Infection Medicine (IMIT) and Institute for Bioinformatics and Medical Informatics (IBMI), University of Tübingen, Tübingen, Germany; Center for Plant Molecular Biology (ZMBP), Interfaculty Institute of Microbiology and Infection Medicine (IMIT), University of Tübingen, Tübingen, Germany; Cluster of Excellence Controlling Microbes to Fight Infections (CMFI), University of Tübingen, Tübingen, Germany; German Centre for Infection Research (DZIF), Partner Site Tübingen, Tübingen, Germany; Department of Biochemistry, University of California Riverside, Riverside, USA; Department of Microbial Bioactive Compounds, Interfaculty Institute of Microbiology and Infection Medicine (IMIT), University of Tübingen, Tübingen, Germany

**Keywords:** synthetic communities, plant leaf microbiomes, pyoverdines, pseudobactin, microbe-microbe interactions, correlation networks, *Arabidopsis thaliana*, secondary metabolites

## Abstract

Microbial communities that promote plant growth show promise in reducing the impacts of climate change on plant health and productivity. Understanding microbe-microbe interactions in a community context is paramount for designing effective microbial consortia that enhance plant resilience. In this study, we investigated the dynamics of a synthetic microbial community (SynCom) assembled from *Arabidopsis thaliana* leaves to elucidate factors shaping community composition and stability. We found notable disparities between *in vitro* pairwise interactions and those inferred from correlation networks *in planta*. Our findings suggested that secondary metabolites, particularly antimicrobials, might mediate interactions *in vitro*, but fade into the background in the community context. Through co-cultivation experiments, we identified the siderophore pseudobactin as a potent antimicrobial agent against several SynCom members, but its impact on community composition *in planta* was negligible. Notably, dominant SynCom members, such as *Pseudomonas koreensis, Flavobacterium pectinovorum*, and *Sporobolomyces roseus*, exhibited only positive correlations, suggesting synergism based on for example exopolysaccharides and biotransformation might drive community dynamics rather than competition. Two correlations between SynCom members in the co-abundance network corresponded with their pairwise *in vitro* interactions, highlighting the potential for further research, and demonstrating the usefulness of correlation networks in identifying key microbe-microbe interactions. Our findings highlight the importance of considering microbiome-wide interaction studies and synthetic communities in understanding and manipulating plant microbiomes.

## Introduction

The microbiome is essential for plant survival: not only does the microbiome promote plant growth, but it also increases stress tolerance to drought, salinity and iron limitation, as well as resistance to pathogens [1–4]. Strategies fighting against climate change to promote plant growth and stress tolerance are becoming more urgent. Engineering the plant microbiome using nature-derived synthetic communities, biocontrol organisms and probiotics can be a prudent way to promote plant growth under the challenging conditions of climate change [2, 3, 5, 6]. As a proof of concept, Schmitz *et al.,* used a synthetic community assembled from the rhizosphere of the desert plant *Indigofera* and were able to increase the salt tolerance of tomato plants [5].

For sustainable and long-term use of synthetic communities as biocontrol agents, understanding the mechanisms that shape and stabilize such communities on plants is crucial [7–9]. In this respect, not only microbe-host interactions but also overlooked microbe-microbe interactions play a role in community dynamics. Correlation networks based on co-abundance and co-occurrence of microbiome members of plants are a promising source for the detection of microbe-microbe dependencies in a community context [10–12]. Correlation network analyses of whole *Arabidopsis thaliana* microbiomes have already shown that microbe-microbe interactions are affected by environmental impacts and the plant phenotype [13–15]. Nevertheless, these factors explain only a part of the dynamics that drive microbiomes. *In vitro* pairwise interactions studies and *in situ* genome mining have revealed the enormous potential of microbiome members to produce secondary metabolites [16–19]. The identification of many genes dedicated to non-ribosomal peptides (NRPs), polyketides (PKs), ribosomally synthesized and post-translationally modified peptides (RiPPs), and toxins in plant microbiomes indicates a rich repertoire of potential antimicrobial agents [17, 20, 21]. Therefore, these metabolites are assumed to play a major role in plant microbiomes, but little is known about how direct pairwise interactions of microbial members are reflected in complex microbiome interactions.

The objective of this study was to elucidate microbe-microbe interactions among core microbiome members of the *A. thaliana* leaf microbiome. As a model system, we used a synthetic community from *A. thaliana* leaves based on high occurrences across multiple plant samples [13], (Chaudhry et al., in preparation). We investigated correlations within the community and with other members of the *A. thaliana* epiphytic microbiome based on co-abundance. We then explored correlations of these microbiome members through *in vitro* pairwise interaction studies and observed significant differences between *in vitro* relations and *in planta* correlation networks. The high number of inhibitions in pairwise interactions suggests that pairwise interactions might be driven by the production of antimicrobial secondary metabolites. This initial observation led us to question why interactions based on antimicrobial compounds are not displayed in correlation networks. By using the siderophore pseudobactin from *Pseudomonas koreensis* as an example, we showed that this strong antimicrobial agent is potent in pairwise interactions but has no effect on the SynCom composition. Strong pairwise inhibitors like *P. koreensis* and *Sporobolomyces roseus* showed a lot of positive correlations *in planta* indicating that competition based on antimicrobials might play a subordinate role in the *A*. *thaliana* leaf microbiome. Our findings help to understand the dynamics within plant-associated microbiomes and highlight microbiome-wide correlation networks and synthetic communities as promising tools for the pre-selection of relevant microbe-microbe interactions in plant microbiome engineering efforts.

## Materials and Methods

### Microbial strains and SynCom assembly

Microbial strains for the construction of a synthetic leaf-associated community from *Arabidopsis thaliana* (SynCom) were isolated through a three years garden experiment by Almario *et al*., Strain selection was based on high occurrence of operational taxonomic units (OTUs) across all plant samples from different seasons (occurrence in ≥ 95% of samples for fungi and ≥ 98 % of samples for bacteria, cut off ≥ 10 reads per sample) as obtained by 16S rRNA / ITS2 MiSeq Illumina amplicon sequencing [13]. Taxonomical classification of SynCom members, comprising 13 bacteria and three fungi (Table 1), was performed through 16S rRNA and ITS2 analysis using Blast. (Chaudhry et al., in preparation)

**Table 1:** SynCom members characterized by 16S rRNA and ITS2 similarity (BlastN)

| Closest Type species match | Short name | Closest type strain match | Percent identity with type strain % |
| --- | --- | --- | --- |
| *Aeromicrobium fastidiosum* | *A. fastidiosum* | DSM 10552(T) | 99.30 |
| *Arthrobacter humicola* | *A. humicola* | KV-653(T) | 100.00 |
| *Bacillus altitudinis* | *B. altitudinis* | 41KF2b(T) | 100.00 |
| *Dioszegia hungarica* | *D. hungarica* | CBS 4214 | 100.00 |
| *Flavobacterium pectinovorum* | *F. pectinovorum* | DSM 6368(T) | 98.61 |
| *Frigoribacterium faeni* | *F. faeni* | 801(T) | 99.82 |
| *Massilia aurea* | *M. aurea* | AP13T | 100.00 |
| *Methylobacterium goesingense* | *M. goesingense* | iEII3(T) | 99.43 |
| *Microbacterium proteolyticum* | *M. proteolyticum* | RZ36(T) | 99.29 |
| *Nocardioides cavernae* | *N. cavernae* | YIM A1136(T) | 99.23 |
| *Paenibacillus amylolyticus* | *P. amylolyticus* | NBRC 15957(T) | 99.49 |
| *Pseudomonas koreensis* | *P. koreensis* | Ps 9-14(T) | 100.00 |
| *Rhizobium skierniewicense* | *R. skierniewicense* | Ch11(T) | 99.64 |
| *Rhodotorula kratochvilovae* | *R. kratochvilovae* | CBS 7436 | 99.82 |
| *Sphingomonas faeni* | *S. faeni* | MA-olki(T) | 99.50 |
| *Sporobolomyces roseus* | *S. roseus* | CBS 486 | 99.29 |

### Culture media and conditions

Bacterial strains were pre-cultured on nutrient agar NA (BD, USA) or in nutrient broth NB (BD, USA) for 48 h. Fungi were pre-cultured on potato dextrose agar PDA (Carl Roth, Germany) potato dextrose broth PDB (Carl Roth, Germany) for 48 h. For cross-streaking experiments with *P. koreensis* WT and mutant, siderophore promotive F-base agar (Merck, Germany) was used. Growth measurements were performed in MM9 minimal medium [22] and enriched MM9 medium, where MM9 was mixed 1:5 with NB, for bacteria and in PDB for fungi (Table S1 and S2). MM9-7 agar was used to culture the SynCom for 5 days for amplicon sequencing (Table S3). All cultures were incubated at 22 °C and liquid cultures were shaken at 120 rpm.

### Sterile plants and plant spraying

Seeds of *A. thaliana* Ws-0 (Wassilewskija) were sterilized over night with chlorine gas. Therefore, seeds were incubated in presence of 4 ml concentrated hydrochloric acid in 100 ml sodium hypo chloride and 35 mbar vacuum. Sterile seeds were immediately sown on 0.5 x MS agar (1.5 mM CaCl_2_, 0.63 mM KH_2_PO_4_, 9.4 mM KNO_3_, 0.75 mM MgSO_4_, 10.3 mM NH_4_NO_3_, 0.55 mM CoCl_2_ x 6 H_2_O, 0.5 mM CuSO_4_ x 5 H_2_O, 50 mM FeNaEDTA, 50.1 mM H_3_BO_3_, 2.5 mM KI, 50 mM MnSO_4_ x H_2_O, 0.5 mM Na_2_MoO_4_ x 2 H_2_O, 15 mM ZnSO_4_ x 7 H_2_O, 8 g/L agar-agar) and grown in a short day chamber (8 h light, 16 h dark, 22 °C) for 1 - 2 weeks. Seedlings were picked and placed in 12 well plates containing 0.5 MS agar. Plants were further incubated for 2 weeks in a short-day chamber.

For spraying, each SynCom strain was pre-cultured in 20 ml of liquid medium. Cultures were harvested after 48 h of incubation through centrifugation at 7.000 rpm for 5 minutes. Afterwards, cells were washed twice and resuspended in 10 ml of 10 mM MgCl_2_. For each strain, the optical density at wavelength 600 nm (OD_600_) was measured and the cultures were diluted to OD_600_ = 0.2. Equal volumes of each strain dilution were combined to form the different SynCom groups used for amplicon sequencing. The mixtures, augmented with 0.02 % silwet 700 for finer droplet distribution, were sprayed onto sterile 3-week-old *A. thaliana* plants using an airbrush system with 2 mbar pressure applied through 2 brushes. Following inoculation plants were incubated in short-day chambers (8 hours light, 16 hours dark) at 22°C.

### Correlation network analysis

Microbial co-abundance networks were performed as described by Mahmoudi *et al*., [23] on OTU tables collected from the leaf microbiome of wild *A. thaliana* samples. In brief, bacterial and eukaryotic (fungal and non-fungal) OTU tables were filtered to retain only those OTUs present in at least 5 samples with more than 10 reads. The OTU tables were used to calculate SparCC correlations [24] (with default parameters) in the FastSpar platform [25]. Permuted P-values for each correlation were derived from 1.000 bootstraps datasets. Only correlations with *P* ≤ 0.001 were kept for further analysis. The preparation of OUT tables from the raw data followed the workflow of Mahmoudi *et al.,* afterwards correlations were calculated as shown in the workflow stored at zenodo repository (Strong pairwise interactions do not drive interactions in a plant leaf associated microbial community), [https://zenodo.org/uploads/11122216]. Cytoscape (version 3.10.0) [26] was used for visualization of interactions of predators and remaining SynCom microorganisms on genus level.Cytoscape (version 3.10.0) [26] was used for visualization of interactions of predators and remaining SynCom microorganisms on genus level.

### Cross streaking experiments

Pairwise interactions of SynCom members were observed on NB and PDA. Solid pre-cultures were taken with cotton swaps, resuspended in 10 mM MgCl_2_ and streaked out on NA/PDA agar plates. Once the test strain was dry, all SynCom members were streaked crosswise onto the test strain. Inhibiting interactions were observed after 48 h incubation by the production of inhibition zones. Promoting interactions were observed by higher growth in contact zones. Cross streaking experiments to test the effect of pseudobactin were performed on F-base agar for 48 h using *P. koreensis* WT and the *ΔpvdI/J* mutant.

### Genome mining

Genomes of SynCom members were analyzed by AntiSMASH 7 [27] for the presence of biosynthetic gene clusters of secondary metabolites. Similarities to known compounds were further investigated by MiBiG [28] comparison and BlastN/BlastP analysis.

### Pseudobactin identification and purification

*P. koreensis* cultivated in 1 L MM9 medium for 48 h at RT and 100 rpm shaking was used for HPLC-MS and NMR analysis. Cells were harvested by centrifugation at 8.000 rpm for 5 minutes and supernatant was collected. The supernatant was tested for the presence of pseudobactin under UV light (365 nm) and by HPLC-MS analysis. HPLC-MS measurements were performed on an Agilent 1260 Infinity (Agilent technology, USA) using a Kinetex 5 µm 100 Å, 100 x 4.6 mm C18 column and a single-quadrupole G6125B MSD in positive ion mode. Analytical HPLC was performed by using the following parameters: 5 µL injection; solvent A: H_2_O (0.1 % TFA); B: acetonitrile (0.1 % TFA); gradient eluent: 10-100% B over 10 minutes, 100 % B for 2 minutes, and requilibration to initial conditions over 3 minutes; flow rate: 1.0 mL/min; UV detection: 254 nm; retention time: 1.2 min; pseudomolecular ion: *m/z* [M+H]+ = 989.4. For the purification of pseudobactin, the supernatant was loaded onto a C18 cartridge (Supleco, USA). The cartridge was washed with 100% water (0.1 % TFA), and pseudobactin was eluted with 10 % acetonitrile (0.1 % TFA). The fraction was dried using a rotary evaporator and lyocell vacuum evaporator. 20 mg of the dry sample was resuspended in methanol (1 mL), and preparative HPLC was performed by using the following parameters: solvent A: water (0.1 % TFA); solvent B: methanol (0.1 % TFA); isocratic eluent: 15 % B; flow rate: 10 mL/min; UV detection: 254 nm; retention time: 20.5 min. Fractions containing pseudobactin were collected, dried, and analyzed by NMR spectroscopy. ^1^H NMR and 2D spectra were recorded at 700 MHz in D_2_O (4.79 ppm). ^13^C NMR spectra were recorded at 175 MHz in D_2_O (not referenced).

### Deletion Mutant creation

For the investigation of interaction between SynCom members and pseudobactin, a pseudobactin deletion mutant was constructed. For the deletion, genes *pvdI* and *pvdJ* were chosen [29, 30]. The deletion of the genes was performed as described by [31]. In short, the plasmid pEX18Gm was used as deletion vector and transferred into *P. koreensis* by conjugation with *E. coli* S17-λ as donor. The genes were introduced into the *P. koreensis* genome through a double crossover event and subsequently eliminated under selection pressure on antibiotic plates. The success for the deletion was confirmed by PCR of the deletion region, HPLC-MS analysis and UV measurement.

### Pseudobactin interaction studies

Feeding experiments were performed by growing SynCom members in medium supplemented with *P. koreensis* WT and *ΔpvdI/J* mutant supernatant. Therefore, the supernatant of 1 L *P. koreensis* WT and mutant was collected by culturing the strains in MM9 medium. Cells were harvested at 8.000 rpm for 5 minutes and supernatant was sterilized by filtering (0.2 µm pore size). The optimal growth media were developed as MM9 and enriched MM9 medium supplemented with the sterile supernatant of *P. koreensis* WT or Δ*pvdI/J* mutant. For a detailed receipt, see supplements (Table S1 and S2). Since MM9 is an iron delimited medium, some SynCom strains were not able to grow under these conditions. For these organisms, enriched MM9 medium was used with minimal additions of NB or PDA medium. For the growth curves, each SynCom strain was pre-cultured, washed and diluted to OD_600_ = 0.2 with MM9 or enriched MM9 medium. 1 ml of each dilution was added into one well of a 24-well plate. Experiments were performed in triplicates. Plates were incubated at 22 °C and 100 rpm shaking, and OD_600_ was measured after T0= 0 h; T1 = 16 h; T2 = 18 h; T3 = 20 h; T4 = 22 h; T5 = 24 h; T6 = 40 h; T7 = 42 h with a TECAN 2000 (Tecan, Switzerland) device. For additional growth curves with *A. humicola*, 48-well plates and 800 µl total volume were used. Strain was diluted to OD_600_=0.2 in the well. For complementing the inhibiting effect of pseudobactin, *A. humicola* was further investigated in WT + FeSO_4_ MM9 medium. For complementing the *ΔpvdI/J* mutant, *A. humicola* was cultivated in *pvdI/J* + pure pseudobactin MM9 medium (Table S1 and S2).

### Amplicon sequencing

#### Sample preparation

For investigating the relative abundance of SynCom members *in vitro*, amplicon sequencing from MM9-7 agar plates was performed. In detail, each SynCom member was pre-cultured in 20 ml liquid medium and harvested after 48 h incubation by centrifugation at 7.000 rpm for 5 minutes. Cells were washed twice using 10 ml of 10 mM MgCl_2_ and resuspended in MgCl_2_. OD_600_ = 1 was adjusted, and strains were mixed in equal volumes.1 ml mixture was streaked on MM9-7 agar plates and incubated for 5 days at 22 °C. After incubation, cells were scratched off the agar in bead filled tubes (MP fastDNA spin kit) and immediately frozen in liquid nitrogen.

For investigating the relative composition of the SynCom *in planta*, amplicon sequencing from plants was performed. Therefore, SynCom WT, SynCom mutant and SynCom pseudobactin sprayed plants were picked in bead filled tubes (MP fastDNA spin kit) after 5 and 9 days of incubation. Tubes were immediately frozen in liquid nitrogen and plants were crushed at – 30°C using a Precellyse device (Bertin, France) (2 x 20 s, 6.400 rpm).

#### DNA isolation

DNA for amplicon sequencing was isolated using the MP fastDNA spin (MP biomedicals, Germany) kit for soil according to manufacturer’s instructions. DNA was eluted in 75 µl elution buffer. Concentration was measured by nanodrop.

#### Library preparation

The library preparation was done following the studies of Agler *et al*., and Mayer *et al*., [15, 32]. Shortly, DNA was used to amplify the 16S rRNA region of bacteria and the ITS2 region of fungi by PCR. A second PCR was used to introduce custom-designed, single indexed Illumina sequencing adapters to each sample. The primers used contained blocking regions to limit the amplification of plant chloroplast DNA as described in the study of Mayer *et al*., All libraries were pooled in equimolar concentrations and sent to NCCT/University of Tübingen for MiSeq Illumina sequencing (300 cycles). Primers and Illumina adapters used in this study can be found in the article of Agler *et al*. [15]

#### Data analysis

Quality control and trimming of raw reads were performed using fastp (v0.23.4) [33] with default parameters. The demultiplexed raw reads were denoised using DADA2 [34] truncating left and right reads at the 250th and 200th positions, respectively, based on a manual inspection of quality scores. The taxonomic analysis of the amplicon sequence variants (ASV) was carried out using QIIME2 with sklearn classifier [35] against the SILVA database (v138, 99%) [36] for bacterial sequencing runs and the UNITE database (v0.9, 99%) [37] for fungal sequencing runs. The raw read counts for ASVs were exported from QIIME artefacts and used in further analysis. Any taxa that have less than 1% cumulative mean relative abundance were grouped under the category “Other” in the figures. ASVs assigned to *Chloroplast sp.* and *Penicillium sp.* were excluded from the analysis.

## Results

### SynCom members are mainly positively correlated with the epiphytic microbiome

The SynCom was assembled in garden experiments from *A thaliana* leaves based on the occurrence of the taxonomic unit in plant samples. Bacteria present in > 98 % and fungi present in > 95 % of plant samples during different seasons were collected [13].

We first analyzed the abundance and connectivity of SynCom members with each other and with other members of the *A. thaliana* microbiome in an extended data set. The data, spanning a 5-year sampling period of wild *A. thaliana* plants across different seasons, was generated by the analysis of relative operational taxonomic unit (OTU) abundance using metagenomic amplicon sequencing of the leaf microbiome [23]. The positive and negative correlations shown in the network were based on co-abundance in all field samples. OTUs showing no significant abundance dependencies (*p* > 0.001) are not shown in the network. Alignment of the most common sequence of each OTU to 16S rRNA and ITS2 sequences of SynCom members facilitated the identification of OTUs closest related SynCom members (Table 2). The generated network (p ≤ 0.001) allowed the identification of positive (blue) and negative (red) correlations between SynCom members and the *A. thaliana* epiphytic microbiome (Fig. 1a).

**Table 2:**
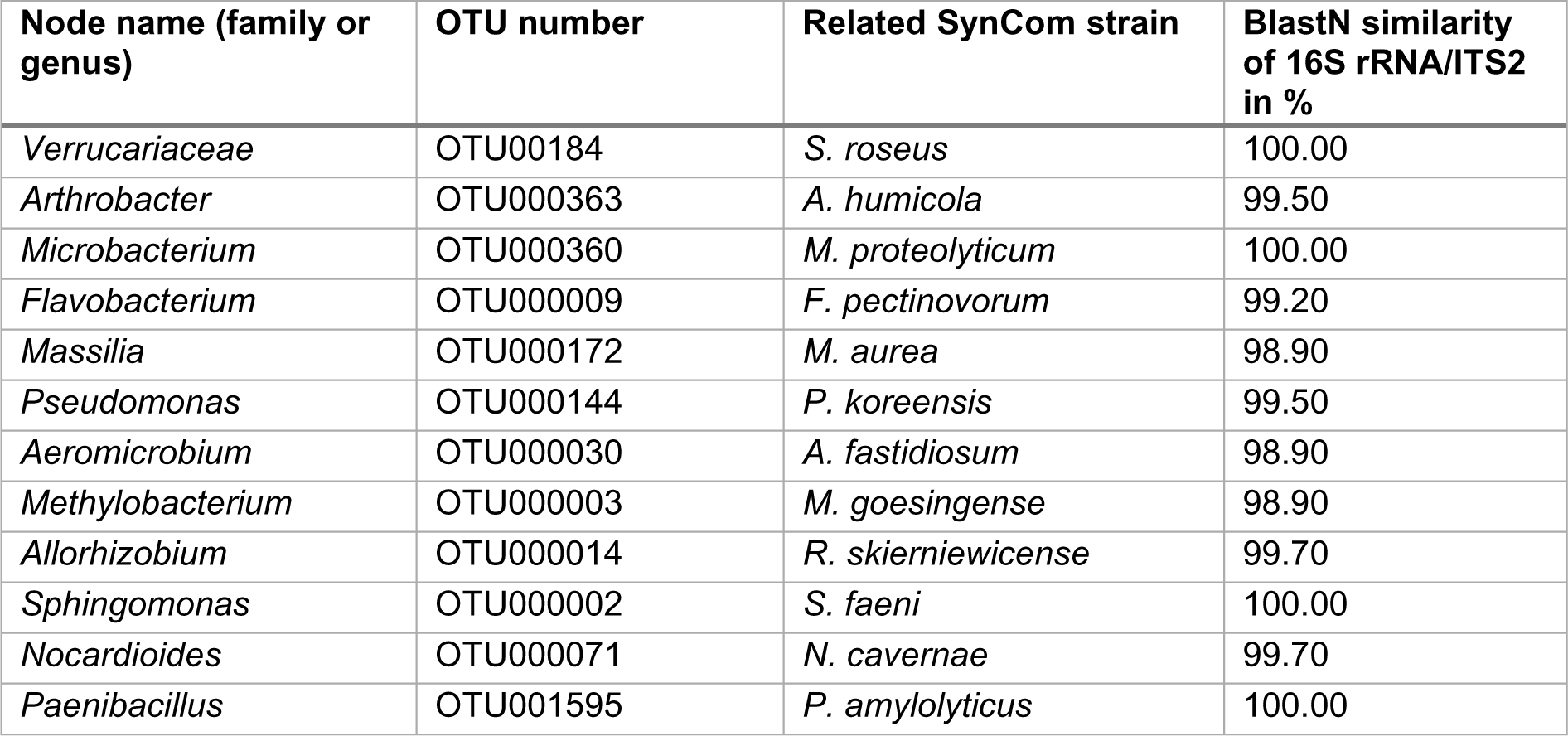
Correlation network OTU annotation by 16S rRNA/ITS2 BlastN against SynCom members. The most common 16S rRNA/ITS2 sequence of each OTU was blasted against 16S RNA/ITS2 regions of SynCom members to identify the closest related nodes in the correlation network for each SynCom members.

**Figure 1:**
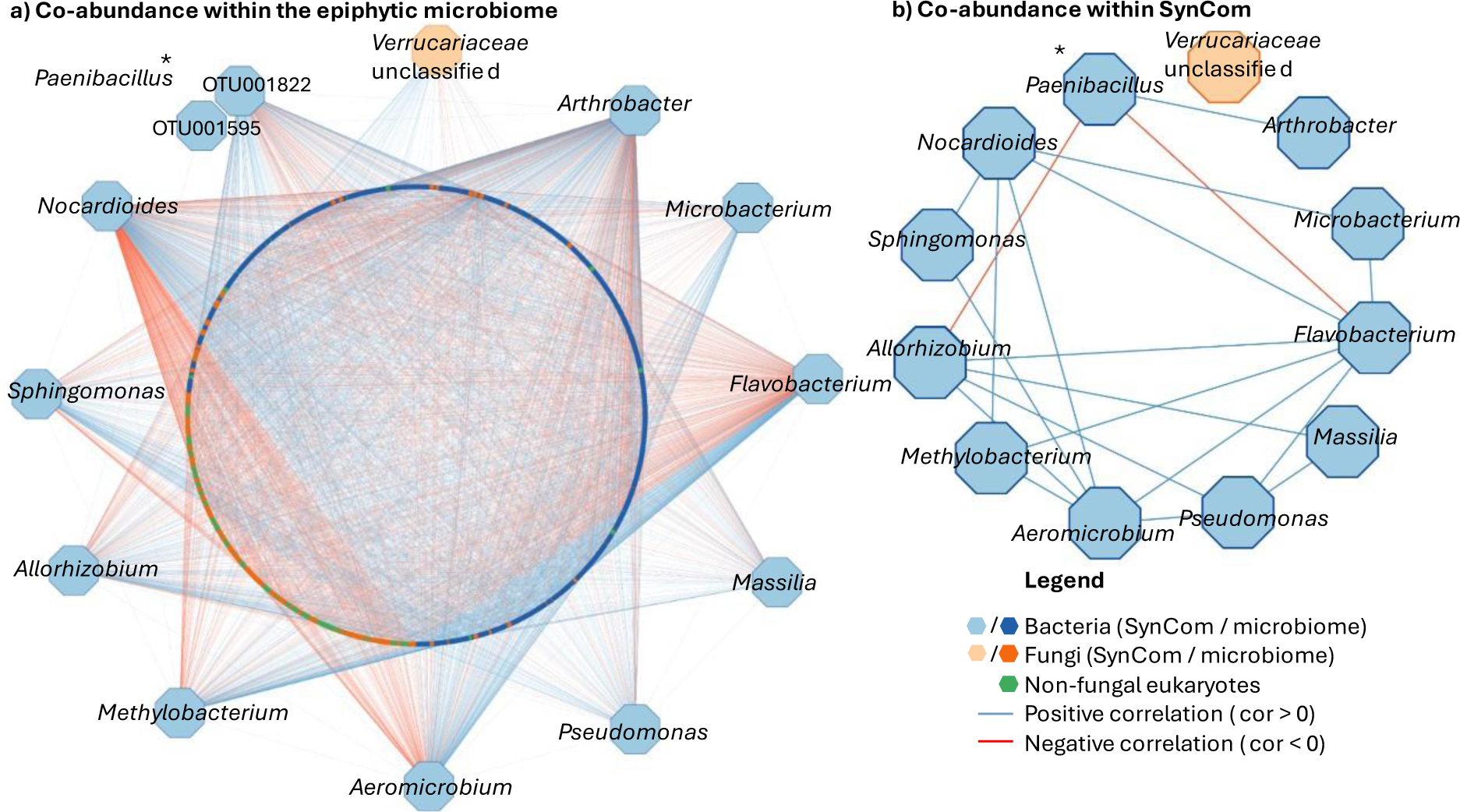
Correlation of SynCom members within the Arabidopsis thaliana epiphytic microbiome based on co-abundance. a) Each node represents an OTU calculated by 16S rRNA/ITS2 Illumina amplicon sequencing. OTUs were identified on genus level. Based on 16S rRNA/ITS2 similarity (BlastN) to SynCom members, closest related OTUs were presented. The network shows correlations with p ≤ 0.001. Positive correlations (cor > 0) are indicated with blue edges, negative correlations (cor < 0) with red edges. * OTU001822 and OTU001595 showed the same BlastN parameters and were both caped in the network. B) SynCom related OTUs and edges were extracted from network a. * OTU001822 showed less, but same correlations as OTU001595 and therefore was replaced.

Overall, SynCom organisms showed a total of 4.116 correlations with OTUs from the phyllosphere microbiome. Among these correlations, the majority (59.5%) was positive, while 40.5% were negative. Four SynCom members, namely *Bacillus altitudinis, Flavobacterium faeni, Dioszegia hungarica*, and *Rhodotorula kratchovilovae*, remained uncorrelated within the network due to their infrequent occurrence and/or low read counts (Table S4). Among the represented SynCom strains, nine exhibited notably high positive correlations (> 60% OTUs positively correlated) within the microbiome. Noteworthy exceptions included *Arthrobacter humicola* (54.6 % OTUs positively correlated), which also displayed the highest total number of correlations (811 interactions), along with *F. pectinovorum* (46.6 % OTUs positively correlated) and *Nocardioides cavernae* (44.1 % OTUs positively correlated). The highest positive correlation was recorded for *S. roseus* (82.8 %), closely followed by *P. koreensis* (74.0 %) (Fig S1). Analysis of connections between SynCom members derived from the microbiome network revealed predominantly positive associations, comprising 20 correlations, with only two relationships exhibiting negative ratios (Fig. 1b). Particularly noteworthy were the highly positive correlations observed for *F. pectinovorum* and *S. faeni*, both displaying positive linkages with six other SynCom members. Notably, *Paenibacillus amylolyticus* emerged as the sole OTU exhibiting negative correlations with two other SynCom members (*F. pectinovorum* and *Rhizobium skierniewicense*). In summary, the SynCom members exhibited predominantly positive correlations within the microbiome, both with other microbiome constituents and among themselves.

### Pairwise interactions do not reflect relations from correlation networks

We further investigated whether relations shown in the correlation network (Fig.1b) could be followed up in pairwise interactions *in vitro*. Therefore, we compared the network data with pairwise interactions observed between SynCom members in cross-streaking experiments on agar plates. Each organism within the SynCom was subjected to cross-streaking against every other member, resulting in a total of 256 tested interactions (Fig 2).

**Figure 2:**
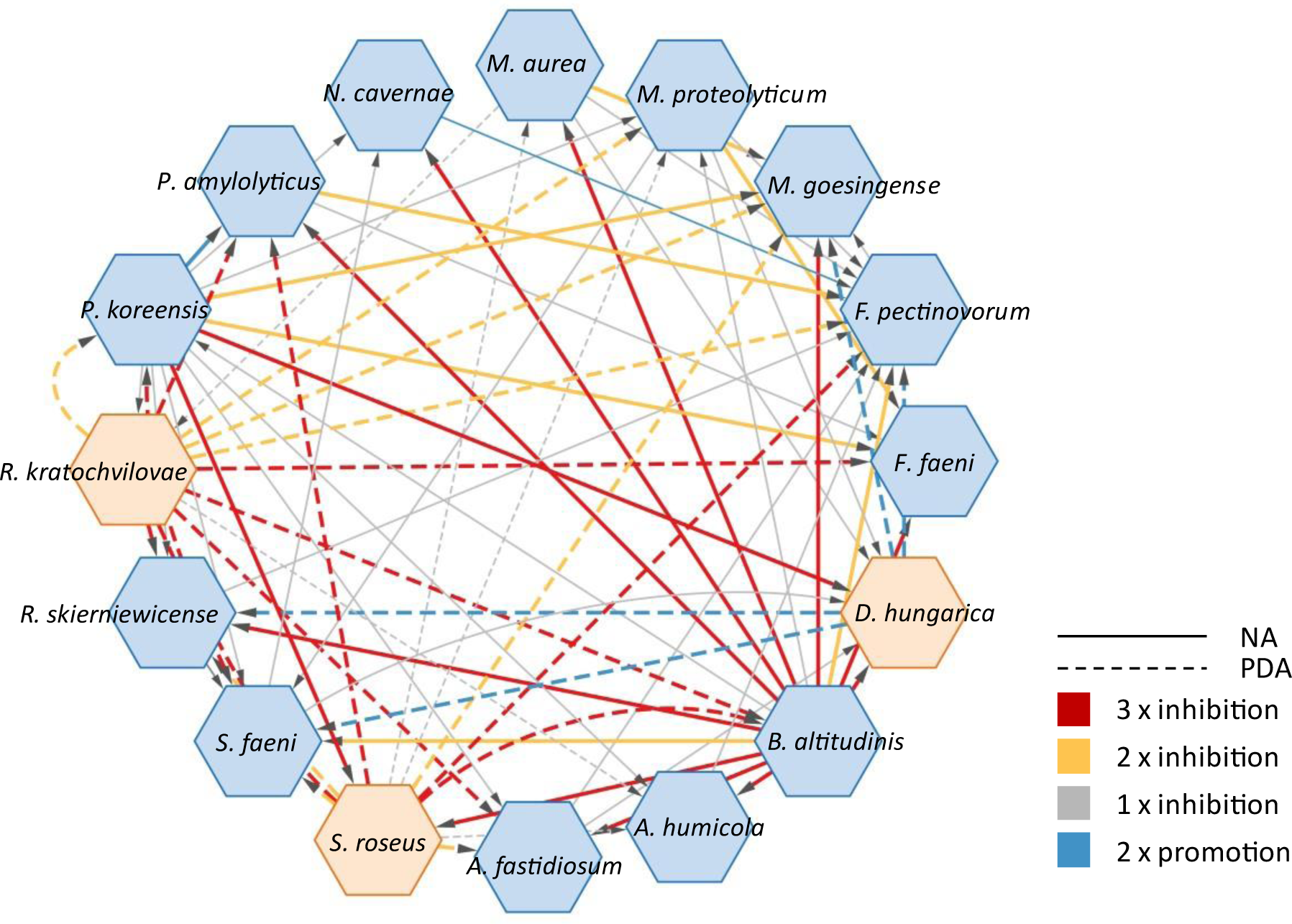
Pairwise interactions of SynCom members in vitro. Strains were grown on optimal growth medium for three days at 22 °C (NA for bacteria, PDA for fungi). Contact zones were assessed for growth-promoting or inhibiting interactions. SynCom bacteria are displayed as blue nodes, SynCom fungi as orange nodes.

While most strains exhibited neutral co-existence, six reproducible growth-promoting interactions were identified among SynCom members across two repetitive experiments. Notably, the yeast *D. hungarica* promoted the growth of four SynCom members (*Methylobacterium goesingense, F. pectinovorum, Sphingomonas faeni*, and *R. skierniewicense*) on their optimal growth agar (PDA). Only one positive interaction was observed between the bacteria *F. pectinovorum* and *N. cavernae*, aligning with the positive correlation observed in the correlation network. The pairwise interaction network predominantly featured negative interactions (Fig. 2). A total of 71 inhibitory relationships were identified, exhibiting 27% reproducibility across three independent experiments, with an additional 24% occurring in two of the three repetitions. Among these, 47 interactions originated from bacteria, and 24 from fungi. Notably, *B. altitudinis* emerged as the most potent bacterial inhibitor within the SynCom, displaying inhibitory effects against 14 SynCom strains in pairwise assessment. However, despite its strong inhibitory activity, *B. altitudinis* was not represented in the correlation networks due to low OTU reads for the strain.

Another prominent inhibitor in pairwise interactions was *P. koreensis*, which inhibited 10 other SynCom members in at least one experiment, with four strains (*M. goesingense, F. faeni, D. hungarica, S. roseus*) being reproducibly inhibited. Interestingly, *P. koreensis* exhibited solely positive correlations with SynCom members within the correlation network.

The most susceptible strain was *F. pectinovorum*, which displayed sensitivity to eight partners in the cross-streaking experiment. However, despite its negative correlation with *P. amylolyticus* in the correlation networks*, F. pectinovorum* showed a high number of positive relationships with other organisms.

Among the fungi, *R. kratochvilovae* (11) and *S. roseus* (10) exhibited the highest number of inhibitory interactions. Whereas *S. roseus* was highly positively correlated in co-abundance networks, it consistently displayed four inhibiting interactions across all pairwise experiments. Interestingly, the inhibitory potential of both fungi was only evident when grown on PDA. *R. kratochvilovae* and *S. roseus* consistently restricted the growth of potent bacterial inhibitors such as *B. altitudinis* and *P. koreensis* on this medium. Conversely, *B. altitudinis* and *P. koreensis* inhibited *S. roseus* on their preferred. These findings underscore the significant impact of optimal nutrient availability on the SynComs pairwise interactions. Collectively, contrary to the correlation network analysis, pairwise interactions unveiled a substantial repertoire of inhibitory interactions among SynCom members. This prompted us to investigate the reasons behind this discrepancy.

### The SynCom encodes a variety of secondary metabolite gene clusters

Antagonistic microbe-microbe interactions are a pervasive phenomenon in pairwise interactions of members from the *A*. *thaliana* leaf microbiome. Most inhibitions are attributed to the vast repertoire of antimicrobial compounds synthesized by a diversity of biosynthetic enzyme classes [16, 17]. To investigate whether the observed inhibitory pairwise interactions are caused by antimicrobial compounds, we analyzed the potential of each SynCom member to produce secondary metabolites. Therefore, we utilized AntiSMASH, a tool for predicting biosynthetic gene clusters (BGCs). Figure 3 illustrates the abundance of BGCs among SynCom strains, totaling 103 gene clusters. *P. amylolyticus* encodes the highest number of BGCs (13), followed by *B. altitudinis* (12), *R. skierniewicense* (11), *P. koreensis* (10), and *M. goesingense* (10). Interestingly, these organisms, except for *M. goesingense,* exhibit significant potential for antimicrobial compound production, based on the presence of RiPP, PKS, NRPS, and hybrid gene clusters. Furthermore, examination of gene clusters from the inhibitor strains in pairwise interactions revealed similarities to BGCs encoding known antimicrobials. For instance, *B. altitudinis* possesses BGCs closely resembling those encoding antimicrobials such as bacilysin (100 % similarity), surfactin (85 % similarity), and bacillibactin (53 % similarity). *P. koreensis* exhibits genes associated with the production of the siderophore pseudobactin from the pyoverdine class, while a 100 % similarity to the BGC of polymyxin B was predicted for one NRPS gene cluster of *P. amylolyticus*. In contrast, strains showing higher sensitivity in pairwise interactions, such as *F. pectinovorum, Frigoribacterium faeni*, and *N. cavernae*, lack NRPS and PKS gene clusters, and are characterized by the presence of terpene and betalactone BGCs (Table S5). The two strong inhibitory fungi *R. kratochvilovae* and *S. roseus* carry a low number of BGCs (4) compared to their bacterial equivalents. Both strains contain two NRPS gene clusters with no similarity to known BGCs. Notably, *D. hungarica* carrying 3 NRPS BGCs shows no inhibition in pairwise interactions.

**Figure 3:**
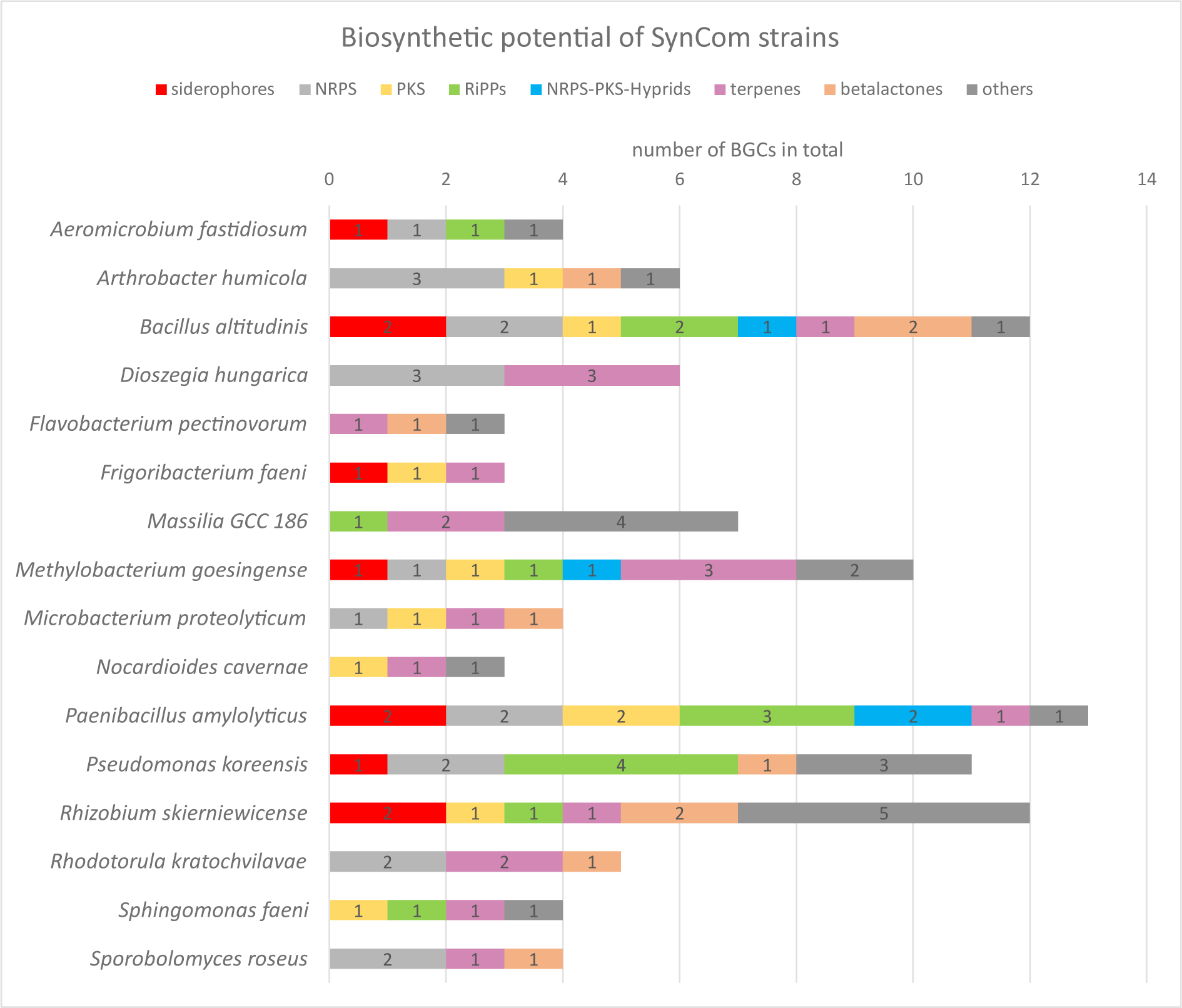
Potential of SynCom strains to produce secondary metabolites. The potential to produce secondary metabolites is based on the presence of biosynthetic gene clusters BGCs as revealed by AntiSMASH 7 analysis.

### Pairwise inhibitors are not inherently dominant strains in the SynCom *in vitro*

As a next step, we wanted to investigate whether pairwise interactions play a role in shaping the SynCom *in vitro*. We posited that inhibitor strains might exhibit a colonization advantage within the community by producing antimicrobial compounds, leading to their dominant abundance. To investigate this, amplicon sequencing of the entire SynCom cultivated together on minimal agar was conducted. Therefore, equal volumes of OD_600_ = 1 mixtures of each strain were mixed. For the experiment, MM9-7 minimal agar was chosen to mimic the limited nutrient bioavailability on plant leaf surfaces [38, 39]. Following a 5-day incubation period, *P. koreensis* emerged as the most prevalent bacterium within the SynCom, with an 89 % relative abundance. Notably, the relative abundance of *P. koreensis* increased fourfold over the incubation period. Regarding fungi, *R. kratochvilovae* showed the highest abundance at 70 %, with a 1.6-fold increase during the growth period (Fig. S3). Interestingly, both *R. kratochvilovae* and *P. koreensis*, recognized as potent inhibitor strains in pairwise interactions, displayed the highest abundance within the SynCom on the plate. In contrast, *B. altitudinis*, able to inhibit 14 SynCom strains in the preceding experiment, showed a ∼ 19-fold reduction in abundance over the incubation period, resulting in a total relative abundance of < 0.5 % (Fig. S2). Strains susceptible to inhibition, such as *F. pectinovorum* and *R. skierniewicense*, were relatively abundant compared to the main bacterial inhibitor strain, *B. altitudinis*. Furthermore, besides *P. koreensis*, *F. pectinovorum* was the only bacterial strain increasing over the incubation period. Although the strong inhibitors *P. koreensis* and *R. kratochvilovae* were dominant colonizers, the low abundance of other strong inhibitor strains like *B. altitudinis* and *S. roseus* indicates that inhibitors do not inherently have a colonization advantage within the SynCom *in vitro.* Therefore, pairwise interactions are not necessarily reflected by the composition of the SynCom on agar plates.

### Pseudobactin drives inhibitory pairwise interactions of *Pseudomonas koreensis*

Next, we aimed to answer the question why strong pairwise interactions are not reflected in the correlation networks. Therefore, we aimed to identify the mechanism behind a specific inhibitory pairwise interaction, track it through subsequent studies from pairwise interactions to co-cultures with the entire SynCom, and finally, examine it *in planta*.

Due to its high abundance in the SynCom on the plate, its ability to inhibit individual SynCom members and its opposing interactions in the correlation network, *P. koreensis* was chosen for further investigations. Previous studies showed that a pyoverdine siderophore with antimicrobial activity contributes to shaping the root microbiome of *A. thaliana*, suggesting its importance in microbial communities [16]. Since a BGC encoding a pyoverdine was detected in the *P. koreensis* genome, we investigated its role in the leaf associated SynCom. The fluorescent compound was isolated, and its structure confirmed by NMR as pseudobactin, a member of the pyoverdine siderophore class (Fig S4-S11). The successful creation of a deletion mutant was verified by HPLC-MS (Fig. S12). Growing strains in the presence or absence of pseudobactin showed inhibitions of growth for eight SynCom members (Fig. 5c) (growth curves of SynCom strains: Fig. S13). *A. humicola,* was significantly inhibited by pseudobactin. The addition iron to *A. humicola* cultures abolished the inhibiting effect (Fig. 5a), leading to normal growth in pseudobactin-containing medium. The restoring of the inhibition by the addition of iron indicates that *P. koreensis* inhibits SynCom members indirectly by the chelation of iron. The addition of purified pseudobactin to the supernatant of the pseudobactin mutant strain, reestablished the inhibiting effect (Fig. 5b).

**Figure 4:**
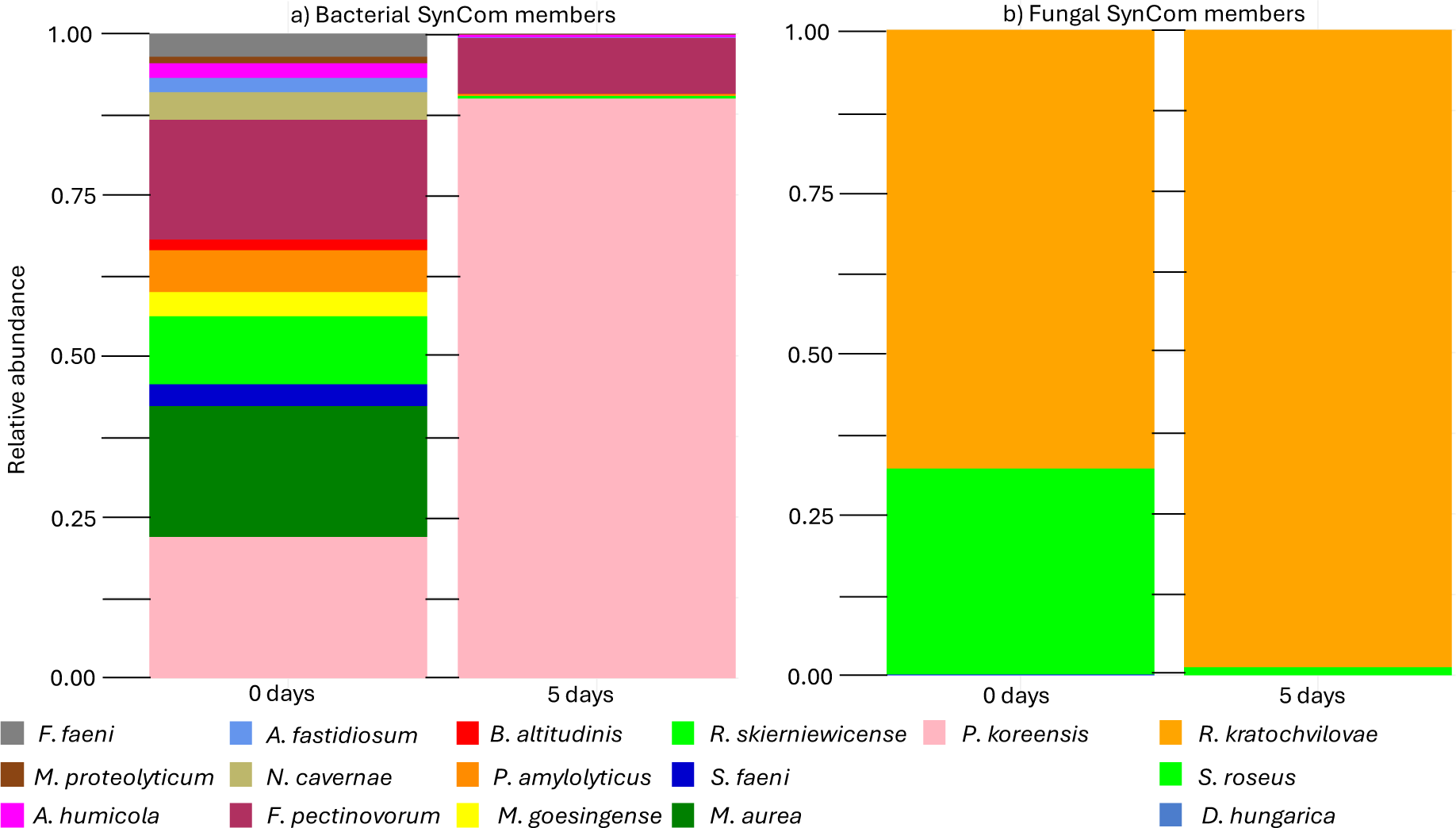
SynCom composition in vitro based on the relative abundance. The SynCom composition *in vitro* was calculated as relative abundance of each strain by 16S rRNA/ITS2 MiSeq Illumina amplicon sequencing from MM9/7 agar at inoculation after 0 days and after 5 days incubation at 22 °C. a) Histograms show the relative abundance of bacterial SynCom members calculated by amplicon sequencing using 16S rRNA specific primers. b) Histograms show the relative abundance of fungal SynCom members calculated by amplicon sequencing using ITS2 specific primers.

**Figure 5:**
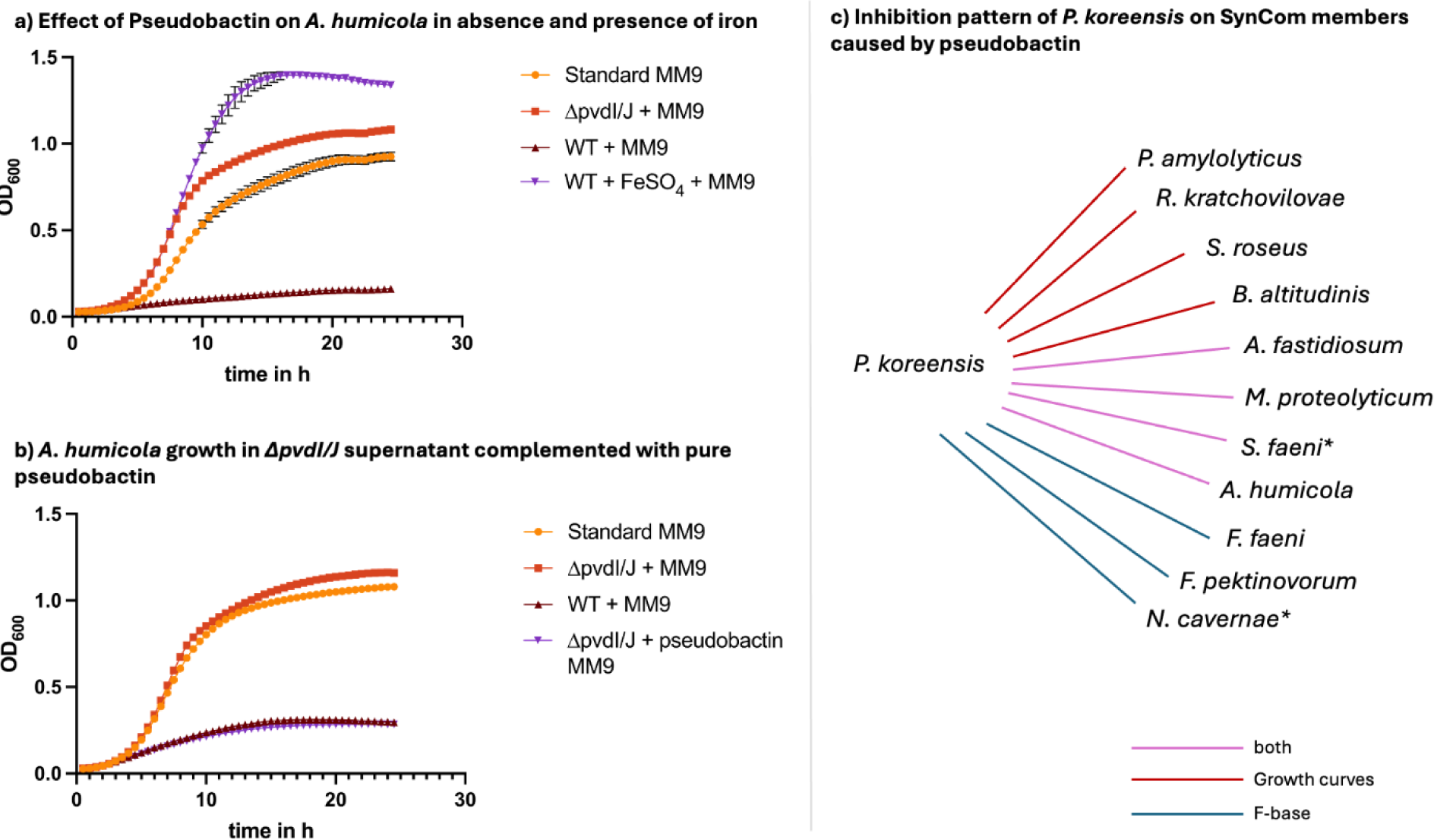
*In vitro* interaction of Pseudobactin with single SynCom members. Growth curves of *A. humicola* were measured automatically in a TECAN 2000 device as OD600 in 1 h intervals at 22 °C and 200 rpm shaking a) Growth curve of *A. humicola* in MM9 medium enriched with sterile supernatant of *P. koreensis* WT and *P. koreensis ΔpvdI/J* mutant. The growth in presence of pseudobactin complemented with FeSo4 is shown in purple. b) Repetition of growth curve of *A. humicola* with complementation of *ΔpvdI/J* mutant supernatant with pure pseudobactin (purple). c) Summary of inhibitions of pseudobactin on SynCom members in growth curves and cross streaking experiment on F-base agar. * Strains showing inhibition zones in contact with both, WT and mutant *P.koreensis* but significantly bigger inhibition zones in contact with WT.

**Figure 6:**
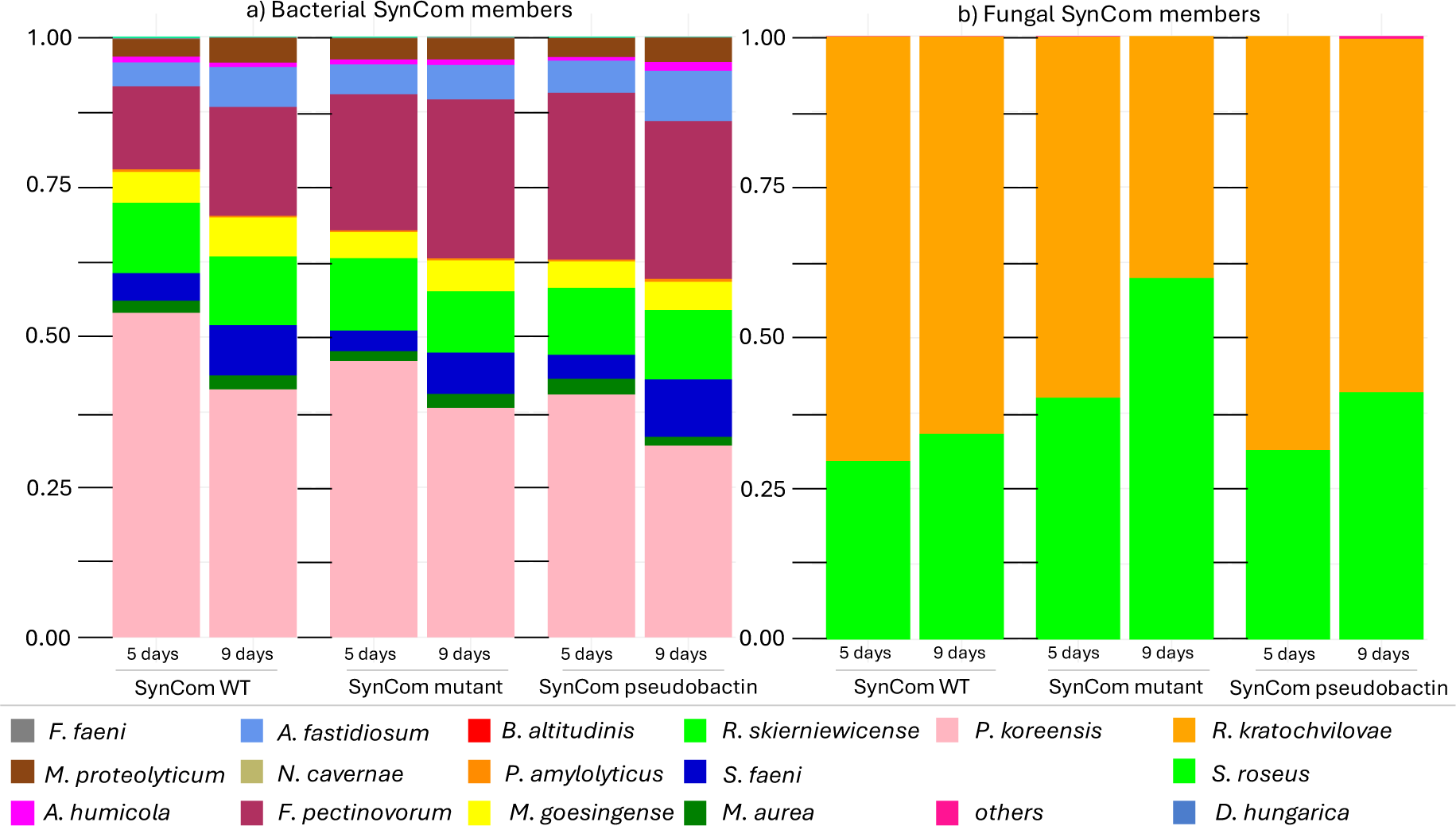
The effect of pseudobactin on the SynCom composition *in planta*. The composition of the SynCom is based on the relative abundance of each member calculated from 16S rRNA/ITS2 MiSeq illumina amplicon data. Three-week-old plants were sprayed with 0.2 OD600 SynCom mixture and incubated at 22 °C in a short light chamber. The sampling was done at two time points (5 days and 9 days after spraying). a) Histograms show the relative abundance of bacterial SynCom members calculated by amplicon sequencing using 16S rRNA specific primers. b) Histograms show the relative abundance of fungal SynCom members (fungi) calculated by amplicon sequencing using ITS2 specific primers.

The inhibitory effects of *P. koreensis* on four SynCom members can therefore be explained by the production of pseudobactin. Notably, three strains, namely *R. kratochvilovae, B. altitudinis*, and *A. humicola*, which were not initially inhibited in pairwise interactions, demonstrated susceptibility when exposed to the siderophore in growth curves. Interestingly, the second most abundant bacterium, *F. pectinovorum*, confirmed its resistance to *P. koreensis* observed in pairwise interactions in the growth curves but showed susceptibility to pseudobactin on F-base agar (Fig. 5c). The remaining sensitive strains from the pairwise interaction study (*N. cavernae*, *M. goesingense, D. hungarica* and *R. skierniewicense)* exhibited instability in growth when cultivated in minimal medium. Consequently, it was not possible to ascertain their growth rate in the presence of pseudobactin, precluding the formulation of definitive statements regarding their response to this antimicrobial agent. Cross streaking experiments of these strains on f-base agar against *P. koreensis* WT and the pseudobactin mutant showed no effect of pseudobactin. Although, *M. goesingense* and *D. hungarica* were sensitive against both, WT and mutant *P. koreensis, they showed larger* zones of inhibition in contact with the wildtype, indicating some sensitivity to pseudobactin but also other compounds produced by P *koreensis.* In summary, pseudobactin is a compound of *P. koreensis* showing antimicrobial activity in pairwise interaction studies on siderophore promotive agar (F-base) and in minimal medium.

### Pseudobactin interactions show no effect in a community context *in planta*

Since the pseudobactin-based inhibitory interactions of *P. koreensis* are not reflected in correlation networks, we further wanted to investigate which role pseudobactin plays within the SynCom *in planta.* Given the known influence of pyoverdines on microbiome composition, we assessed the contribution of pseudobactin by applying three different SynCom preparations to sterile *A. thaliana* plants through plant spraying: the wild type SynCom containing *P. koreensis* (SynCom WT), the SynCom with the P. koreensis pseudobactin mutant (SynCom mutant), and the SynCom mutant supplemented with pure pseudobactin (SynCom pseudobactin). Using amplicon sequencing, we determined SynCom member abundance on plants. After a 5-day and a 9-day incubation period, *P. koreensis* and *F. pectinovorum* emerged as the dominant bacteria, while *R. kratochvilovae* prevailed as the dominant yeast across all three experimental groups. Notably, there were no significant differences in SynCom overall relative abundance between all three groups (SynCom WT, SynCom mutant, SynCom pseudobactin). To assess whether the presence or absence of pseudobactin had an impact on individual SynCom members, the relative abundance of each strain was separately analyzed, revealing no significant alterations among the different groups (Fig. S14). The results display that even though it shows strong inhibiting activity on SynCom strains in pairwise interactions, pseudobactin does not affect the SynCom composition or abundance of any member *in planta*.

## Discussion

In our study we aimed to investigate microbe-microbe interactions of a synthetic plant leaf community to understand which dynamics shape and stabilize the microbiome. Our findings revealed notable disparities between pairwise interactions observed *in vitro* and those inferred from correlation networks *in planta*. Whereas the correlations in the microbiome were mainly positive, pairwise interactions showed a huge number of inhibitory interactions between SynCom members. The huge repertoire of genes to produce secondary metabolites indicated that pairwise interactions are driven by antimicrobial compounds. Accordingly, we identified pseudobactin from *P. koreensis* as a potent antimicrobial agent against several SynCom members in pairwise interaction experiments. However, pseudobactin had no effect on the SynCom composition *in planta*, mirroring the correlation networks, where *P. koreensis* showed no negative correlation to any SynCom member.

### Pairwise interactions do not affect co-abundance of SynCom members in the epiphytic microbiome

Pairwise interaction studies via cross streaking experiments are a common method for the identification of secondary metabolites, especially antimicrobial compounds. AntiSMASH analysis revealed the high potential of *P. koreensis, B. altutidinis* and *P. amylolyticus* to produce secondary metabolites, and it was already shown that secondary metabolites produced by plant microbiome members drive strong pairwise interactions [16] . Therefore, it is most likely that the observed inhibitions in the cross-streaking experiments are based on antimicrobials. Why the interactions within our SynCom are not reflected in the correlation networks remains unclear. Even the addition of pure pseudobactin did not alter the SynCom significantly, suggesting that the inhibitory effect of pseudobactin is limited to pairwise interactions and plays a subsidiary role in a microbiome context. Although Getzke *et. al.* could show an effect of pyoverdine on the composition of the root microbiome, pyoverdines might play a minor role in shaping plant leaf associated communities as compared to root microbiomes. Indeed, while in the rhizosphere iron is limiting and production of siderophores may confer a growth advantage [39, 40] the phyllosphere has higher iron concentrations, decreasing the need for siderophore production [41].

We hypothesized that strong inhibitors would have a colonization advantage in communities. Therefore, we anticipated an increase in the relative abundance of the inhibitory strain and a decrease in that of the sensitive strains over the incubation period. However, for *B. altitudinis*, we observed the opposite effect. Although the genome of this strain encodes a high potential to produce secondary metabolites — possibly a reservoir activated only in the presence of certain competitors or pathogens [42, 43] — the strong inhibitor *B. altitudinis* might be constrained by the community. Long-term co-evolution of plant microbiomes has allowed the adjustment of an optimal balance in microbial composition and microbe-microbe interactions [39, 44, 45]. Thus, it is not surprising that a potent member of the synthetic community (SynCom) is unable to dominate others, despite its substantial antimicrobial potential. This restraint, for example through suppressed production of antimicrobial compounds, can be further investigated using transcriptomic approaches. Tyc *et al*., demonstrated that the antimicrobial activity of certain soil microbiome members is significantly suppressed in co-cultures with commensals compared to monocultures, attributing this to interference with the quorum sensing apparatus or nutrient limitations [46].

Nevertheless, the question remains what *P. koreensis, F. pectinovorum* and *S. roseus* have in common to assert their dominant abundance in the SynCom. It is known that both, *S. roseus* and *P. koreensis*, produce exopolysaccharides (EPS), which help them survive harsh environmental conditions [47]. It is furthermore known that EPS are highly involved in the production of biofilms and that microbiome members can benefit from biofilm producers in their community [48, 49]. *S. roseus* is additionally capable of breaking down leaf surface waxes improving the surface adhesion of organisms on plants [50]. Highlighting their supportive roles, *S. roseus* and *P. koreensis* showed the highest number of positive correlations with the epiphytic microbiome. Their positive linkage strengthens recent findings that adaption by using extracellular metabolites might be a driving force within microbial communities rather than competition by producing antimicrobials [51]. Within the SynCom, *P. koreensis* was again one of the strains counting high number of positive relations but was exceeded by *F. pectinovorum* and *S. faeni*. Interestingly, *F. pectinovorum* was also a dominant bacterium in the SynCom *in vitro* and *in planta*, despite being highly sensitive in pairwise interactions. *Flavobacterium spp*. are common members of plant microbiomes and known for their great ability to degrade extracellular macromolecules like starch. Furthermore *Flavobacterium spp*. indirectly promote plant growth, suggesting that they support the microbiome by biotransformation [10, 52, 53]. The high abundance and positive linkage of *F. pectinovorum* in the SynCom strengthens the hypothesis that synergism plays a huge role in shaping microbial communities.

### Correlation networks for the pre-selection of relevant microbe-microbe interactions

Two correlations of SynCom members in the correlation network mirrored the findings in pairwise interactions. *N. cavernae* and *F. pectinovorum* showed a growth promoting effect in cross-streaking experiments and were positively correlated in the epiphytic microbiome. Moreover, *P. amylolyticus* inhibited *F. pectinovorum in vitro* and the strains were negatively correlated in the microbiome. Whether the pairwise interactions for these strains are truly reflected in correlation networks needs to be further investigated. The mechanistic basis behind the positive connections of *N. cavernae* and *F. pectinovorum* in the plant microbiome is not yet understood. For *P. amylolyticus* strains it was already shown that they are able to produce polymyxin antibiotics [54, 55]. Interestingly, the *P. amylolyticus* strain from the SynCom carries a gene cluster with 100 % similarity to polymyxin B. The production of the compound might explain the antimicrobial activity in pairwise interactions since it is potent against gram negative bacteria like *F. pectinovorum* [56]. Whether the inhibitory interaction is so dominant to be observed within the epiphytic microbiome in correlation networks, remains unknown but it is a promising start for future investigations. Furthermore, it shows that correlation networks are promising methods for preselecting microbe-microbe interactions involved in microbiome shaping. Several publications successfully used bottom-up methods such as pairwise interaction analysis for further investigations in microbial communities and microbiomes [16, 17, 57].

However, our findings indicate that when investigating microbiome interactions on a pairwise basis, there is a high likelihood that these interactions prove to be less significant than expected. As soon as three and more interaction partners exist together in a model system, the complexity of the interaction network increases drastically, limiting the meaningfulness of pairwise interaction approaches [58]. Therefore, beyond-pairwise interaction methods especially computational approaches are getting more and more into the focus of research [59, 60]. Our results demonstrate the limitations of pairwise interaction approaches and suggest the use of microbiome-wide studies like correlation networks for the investigation of dynamics shaping and stabilizing microbial communities. Furthermore, the use of synthetic communities can give insights into the importance of a compound or strain in a microbiome context and therefore is a promising method for investigating microbiome dynamics.

## Data Availability

1. The raw datasets generated from amplicon sequencing for the relative abundance of SynCom members *in vitro* and *in planta* are available in the Zenodo repository (Strong pairwise interactions do not drive interactions in a plant leaf associated microbial community), [https://zenodo.org/records/11122216]
2. The datasets for the visualization of the correlation networks are available in the Zenodo repository (Strong pairwise interactions do not drive interactions in a plant leaf associated microbial community), [https://zenodo.org/records/11122216]
3. The OUT data and workflows used for correlation network calculation are available in the Zenodo repository (Strong pairwise interactions do not drive interactions in a plant leaf associated microbial community), [https://zenodo.org/records/11122216]
4. All raw data for correlation networks based on co-abundance analysed during this study are included in the published article of Mahmoudi *et al*., and its supplementary information files.

## Acknowledgements

NZ, EK, HBO, CH, FH, VC, DP acknowledge funding of the study by the Cluster of Excellence EXC 2124: Controlling Microbes to Fight Infection (CMFI, project ID 390838134). NZ and CB acknowledge funding by the German Federal Ministry for Education and Research (BMBF, Grant MicroMATRIX161L0284C). Furthermore, NZ is grateful for the funding by the German Center for Infection Research (DZIF, Grant TTU09.716). EK, VC, MM and EKl Have been funded by the European Research Council (ERC) under the DeCoCt research program (grant agreement: ERC-2018-COG 820124). PS thanks the European Union’s Horizon Europe research and innovation programm for support through a arie Skłodowska-Curie fellowship n.101108450-MeStaLeM. All authors thank Dr. Libera Lo Presti for helpful comments on the manuscript. Prof. Christoph Mayer for providing the pEXGm18 vector system and Prof. Ewa Musiol-Kroll for *E.coli* S17-λ. We furthermore thank the NGS competence center Tübingen (NCCT) of the University Tübingen for the Illumina MiSeq amplicon sequencing.

## Author contribution

FH, VC, HBO, EK, NZ designed the research; FH conducted the experiments and analyzed the data; CB analyzed data obtained by Illumina amplicon sequencing; MM participated in the analyzation of the data for correlation networks; CH, PS, DP conducted HPLC-MS analyses and structure elucidation. LB created the *ΔpvdI/J* mutant; EKl participated in designing and preparing the library for amplicon sequencing. FH, HBO, EK, NZ wrote the paper.

