## supplemental for "Strong pairwise interactions do not drive interactions in a plant leaf associated microbial community": supplemental_information.docx

**List of Tables**

Table S1: MM9 medium used in growth curves for investigating the effect of pseudobactin on SynCom members. ………………………………………………………………………………………………………...3

Table S2: Enriched MM9 medium used in growth curves for investigating the effect of pseudobactin on SynCom members. …………………………………………………………………………...…………………3

Table S3: MM9/7 defined minimal agar for 16S rRNA/ITS2 amplicon sequencing of SynCom strains in vitro. ……………………………………………………………………………………………………………….4

Table S4: OTUs representing SynCom members which were removed from correlation network. …….4

Table S5: AntiSMASH biosynthetic gene cluster prediction of SynCom strains and similarity to known BGCs. ……………………………………………………………………………………………………..……6,7

**Table S1: MM9 medium used in growth curves for investigating the effect of pseudobactin on SynCom members**. The medium was highly iron-limited and used for growth curves of A. humicola and S. faeni.

| Medium additive | Standard MM9 | WT + MM9 | ∆pvdI/J + MM9 | WT + FeSO4 + MM9 | *∆pvdI/J* + Pseudobactin + MM9 |
| --- | --- | --- | --- | --- | --- |
| MM9 medium | 100 ml | 90 ml | 90 ml | 90 ml | 90 ml |
| WT supernatant | - | 10 ml | - | 10 ml | 10 ml |
| *∆pvdI/J* supernatant | - |  | 10 ml | - | 10 ml |
| FeSO_4_ (1.5 mg/ml) | - |  |  | 100 µl |  |
| Pure Pseudobactin (100 µg/ml) | - |  |  |  | 100 µl |

**Table S2: Enriched MM9 medium used in growth curves for investigating the effect of pseudobactin on SynCom members**. Strains not able to grow in iron-limited MM9 medium were grown in enriched MM9 containing low amounts of optimal growth medium NB (for bacteria) or PDB (for yeasts). Enriched MM9 medium was used for growth curves of B. altitudinis, A. fastidiosum, P. amylolyticus, M. pectinovorum, N. cavernae, M. goesingense, R. skierniewicense, D. hungarica, S. roseus, R. kratchovilovae.

| Medium additive | Standard enriched MM9 | WT + MM9 | ∆pvdI/J + MM9 | WT + FeSO4 + MM9 | *∆pvdI/J* + pseudobactin + MM9 |
| --- | --- | --- | --- | --- | --- |
| MM9 medium | 80 ml | 70 ml | 70 ml | 70 ml | 70 ml |
| NB/PDA | 20 ml | 20 ml | 20 ml | 20 ml | 20 ml |
| WT supernatant | - | 10 ml | - | 10 ml |  |
| *∆pvdI/J* supernatant | - |  | 10 ml | - | 10 ml |
| FeSO_4_ (1.5 mg/ml) | - |  |  | 100 µl |  |
| Pure pseudobactin (100 µg/ml) | - |  |  |  | 100 µl |

**Table S3: MM9/7 defined minimal agar for 16S rRNA/ITS2 amplicon sequencing of SynCom strains in vitro**. MM9 medium was modified as shown in the table to obtain a defined minimal agar suitable for amplicon sequencing and inspired by the plant leaf surface.

| Solution | Chemical | Volume |
| --- | --- | --- |
| Pre autoclave solution in 950 ml | KH_2_PO_4_ | 0.30 g |
|  | NaCl | 0.50 g |
|  | NH_4_Cl | 1.00 g |
|  | Agar | 15.00 g |
| Autoclave at 121 °C, 20 min | | |
| Post autoclave solution (filter sterilize 0.2 µm and add to pre autoclave solution) | Glucose 20 % | 10.00 ml |
|  | MgSO_4_ 1 M | 1.00 ml |
|  | CaCl_2_ 100 mM | 1.00 ml |
|  | amino acid solution | 30.00 ml |
|  | Trace element solution | 10.00 ml |
| Amino acid solution preparation filter sterilized (0.2µm)   - Mix equal volumes of each solution I-VI | | |
| Solution I (in 100 ml dH_2_O) | Phe | 0.99 g |
|  | Lys | 1.10 g |
|  | Arg | 2.50 g |
| Solution II (in 100 ml dH_2_O) | Gly | 0.20 g |
|  | Val | 0.70 g |
|  | Ala | 0.84 g |
|  | Trp | 0.41 g, |
| Solution III (in 100 ml dH_2_O) | Thr | 0.71g |
|  | Ser | 8.40 g |
|  | Pro | 4.60 g |
|  | Asn | 0.96 g |
| Solution IV (in 90 ml dH_2_O + 10 ml HCl (36 %)) | Asp (free acid) | 1.04 g |
|  | Gln | 14.60 g |
| Solution V (dissolve K.Glu in 80 ml dH_2_O, add rest and fill up to 100 ml with dH_2_O) | K.Glu | 18.70 g |
|  | Tyr | 0.36 g |
|  | NaOH | 4.00 g |
| Solution VI (in 100 ml dH_2_O) | Ile | 0.79 g |
|  | Leu | 0.77 g |
| Trace element solution preparation (filter sterilized 0.2 µm) | | |
| EDTA-solution (in 800 ml dH_2_O, pH 7.5) | EDTA | 5.00 g |
| Final solution (fill up to 1 L with dH_2_O) | FeCl_3_ - 6 H_2_O | 0.83 g |
|  | ZnCl_2_ | 84.00 mg |
|  | CuCl_2_ - 2H_2_O | 13.00 mg |
|  | CoCl_2_ - 2H_2_O | 10.00 mg |
|  | H_3_BO_3_ | 10.00 mg |
|  | MnCl_2_ - 4H_2_O | 1.60 mg |

**Table S4: OTUs representing SynCom members which were removed from correlation network.** The OTUs showed highest BlastN similarity to the 16S rRNA/ITS2 sequence of the named SynCom member. OTUs showing < 10 reads per sample and/or occurrence in < 5 samples were removed.

| **OTU** | **related SynCom strain** | **BlastN similarity** | **samples with OTU occurence** | **samples with > 10 reads** |
| --- | --- | --- | --- | --- |
| Otu002983 | *B. altitudinis* | 98.70% | 9 | 3 |
| Otu004835 | *F. faeni* | 100.00% | 35 | 0 |
| Otu02956 | *D. hungarica* | 100.00% | 29 | 0 |
| Otu00955 | *R. kratchovilovae* | 100.00% | 3 | 2 |

Number of OTUs correlated to SynCom members within the Arabidopsis thaliana epiphytic microbiome

|  | #REF! | posititive cor total | negative cor total | positive cor syncom | negative cor syncom |
| --- | --- | --- | --- | --- | --- |
| S. roseus | 1 | 53 | 11 | 0 | 0 |
| A. humicola |  | 519 | 292 | 1 | 0 |
| M. proteolyticum |  | 110 | 48 | 2 | 0 |
| F. pectinovorum |  | 255 | 292 | 6 | 1 |
| M. aurea |  | 98 | 44 | 2 | 0 |
| P. koreensis |  | 111 | 39 | 4 | 0 |
| A. fastidiosum |  | 215 | 179 | 6 | 0 |
| M. goesingense |  | 199 | 104 | 3 | 0 |
| R. skiernervicense |  | 178 | 85 | 4 | 1 |
| S. faeni |  | 155 | 71 | 2 | 0 |
| N. cavernae |  | 309 | 392 | 5 | 0 |
| P. amylolyticus |  | 246 | 111 | 1 | 2 |

**Figure S1: Total number of OTUs connected to SynCom members by edges in correlation networks based on co-abundance**. Total positive correlations (cor > 0) (light blue) and negative correlations (cor < 0) (red) of SynCom members to the epiphytic microbiome of A. thaliana. Positive (dark blue) and negative (organe) correlations of SynCom members to each other extracted of the whole correlation network.

**Table S5: AntiSMASH biosynthetic gene cluster prediction of SynCom strains and similarity to known BGCs.** AntiSMASH 7.0 was used for the identification of BGCs of SynCom members and their similarity to known clusters. *Two split NRPS BGCs of P. koreensis were involved in the production ofpseudobactin and counted as one cluster in Fig. 3

| **Strain** | **Cluster prediction** | **Most similar known cluster** | **similarity** |
| --- | --- | --- | --- |
| A. *fastidiosum* | redox-cofactor |  |  |
|  | NI-sierophore | Desferroxamine E | 75 % |
|  | RiPP-like |  |  |
|  | NAPAA | e-Poly-L-Lysine | 100 % |
| A. *humicola* | NRPS-like | SLI- 2138 | 11 % |
|  | Type 3 PKS | pentalenolactone | 15 % |
|  | betalactone | microansamycin | 7 % |
|  | NAPAA | stenothricin | 31 % |
|  | NAPAA |  |  |
|  | RRE-containing |  |  |
| B. *altitudinis* | betalactone | - | - |
|  | RiPP-like |  |  |
|  | Type 3 PKS |  |  |
|  | NRPS | lichenysin | 85 % |
|  | NRP-metallophore | bacillibactin | 80 % |
|  | RiPP-like |  |  |
|  | Type 1 PKS / NRPS | Zwittermycin A | 18 % |
|  | betalactone | fengycine | 53 % |
|  | Terpene |  |  |
|  | NRPS-like | locillomycin | 21 % |
|  | NI-siderophore | schizokinen | 60 % |
|  | RRE-containing |  |  |
| *F. pectinovorum* | Arylpolyene/resorcinol | flexirubin | 91 % |
|  | terpene | carotenoid | 28 % |
|  | betalactone |  |  |
| *M. aurea* | terpene |  |  |
|  | RiPP-like | paulomycin | 3 % |
|  | arylpolyene | APE Vf | 35 % |
|  | hserlactone |  |  |
|  | Hserlactone, RRe-containing |  |  |
|  | Hydrogen-cyanide |  |  |
|  | terpene | carotenoid | 100 % |
| *M. goesingense* | terpene | carotenoid | 100 % |
|  | RiPP-like |  |  |
|  | Redox-cofactor |  |  |
|  | Type 1 PKS | Oryzanaphthopyran A | 6 % |
|  | NRP-metallophore | taiwachelin | 22 % |
|  | Hserlactone |  |  |
|  | Terpene |  |  |
|  | Terpene |  |  |
|  | Type 1 PKS/ NRPS |  |  |
|  | NAPAA |  |  |
| *M. proteolyticum* | betalactone | microansamycin | 7 % |
|  | terpene | carotenoid | 21 % |
|  | NAPAA | e-Poly-L-Lysin | 100 % |
|  | Type 3 PKS |  |  |
| *N. cavernae* | terpene | carotenoid | 14 % |
|  | Betalactone / NRPS-like | Formicamycins A-M | 4 % |
|  | Type 3 PKS | alkylresorcinol | 100 % |
| *P. amylolyticus* | Type 3 PKS |  |  |
|  | Type 3 PKS | Corynecin II | 13 % |
|  | NRPS-like |  |  |
|  | lassopeptide | paeninodin | 60 % |
|  | proteusin |  |  |
|  | NI-siderophore |  |  |
|  | Trans-AT PKS / NRPS | pellasoren | 33 % |
|  | Trans-AT PKS / NRPS | paenilipoheptin | 23 % |
|  | terpene | carotenoid | 33 % |
|  | Opine-like-metallophore | bacillopaline | 100 % |
|  | Lanthipeptide-class-ii | Gramicidin S | 15 % |
|  | Lanthipeptide-class-iv |  |  |
|  | NRPS | polymyxin | 100 % |
| *P. koreensis* | NAGGN |  |  |
|  | NRPS | Pf-5 pyoverdine* | 21 % |
|  | arylpolyene | APE Vf | 40 % |
|  | NRPS-like | fragin | 37 % |
|  | RiPP-like |  |  |
|  | NRP-metallophore | Pf-5 pyoverdine* | 10 % |
|  | RiPP-like |  |  |
|  | RiPP-like |  |  |
|  | betalactone | fengycin | 13 % |
|  | Hydrogen-cyanide | Hydrogen cyanide | 100 % |
|  | Redox-cofactor | Lankacidin C | 13 % |
|  | RiPP-like |  |  |
| *R. skierniewicense* | terpene |  |  |
|  | arylpolyene | Persiamycin A | 5 % |
|  | Lanthipeptide-class V |  |  |
|  | NI-siderophore | Desferrioxamine E | 50 % |
|  | betalactone | xantholipin | 4 % |
|  | NI-siderophore | roseobactin | 50 % |
|  | thioamitides |  |  |
|  | betalactone |  |  |
|  | hserlactone |  |  |
|  | Hydrogen-cyanide |  |  |
|  | hserlactone |  |  |
|  | Type 1 PKS |  |  |
| *S. faeni* | RiPP-like |  |  |
|  | terpene | carotenoid | 50 % |
|  | Type 3 PKS |  |  |
|  | Redox-cofactor | Lankacidin C | 13 % |
| D. *hungarica* | Terpene |  |  |
|  | NRPS-like |  |  |
|  | NRPS-like |  |  |
|  | terpene |  |  |
|  | NRPS-like |  |  |
|  | terpene |  |  |
| *R. kratchovilovae* | NRPS-like |  |  |
|  | NRPS |  |  |
|  | Terpene |  |  |
|  | Betalactone |  |  |
|  | Terpene |  |  |
| *S. roseus* | NRPS-like |  |  |
|  | Terpene |  |  |
|  | Betalactone |  |  |
|  | NRPS |  |  |

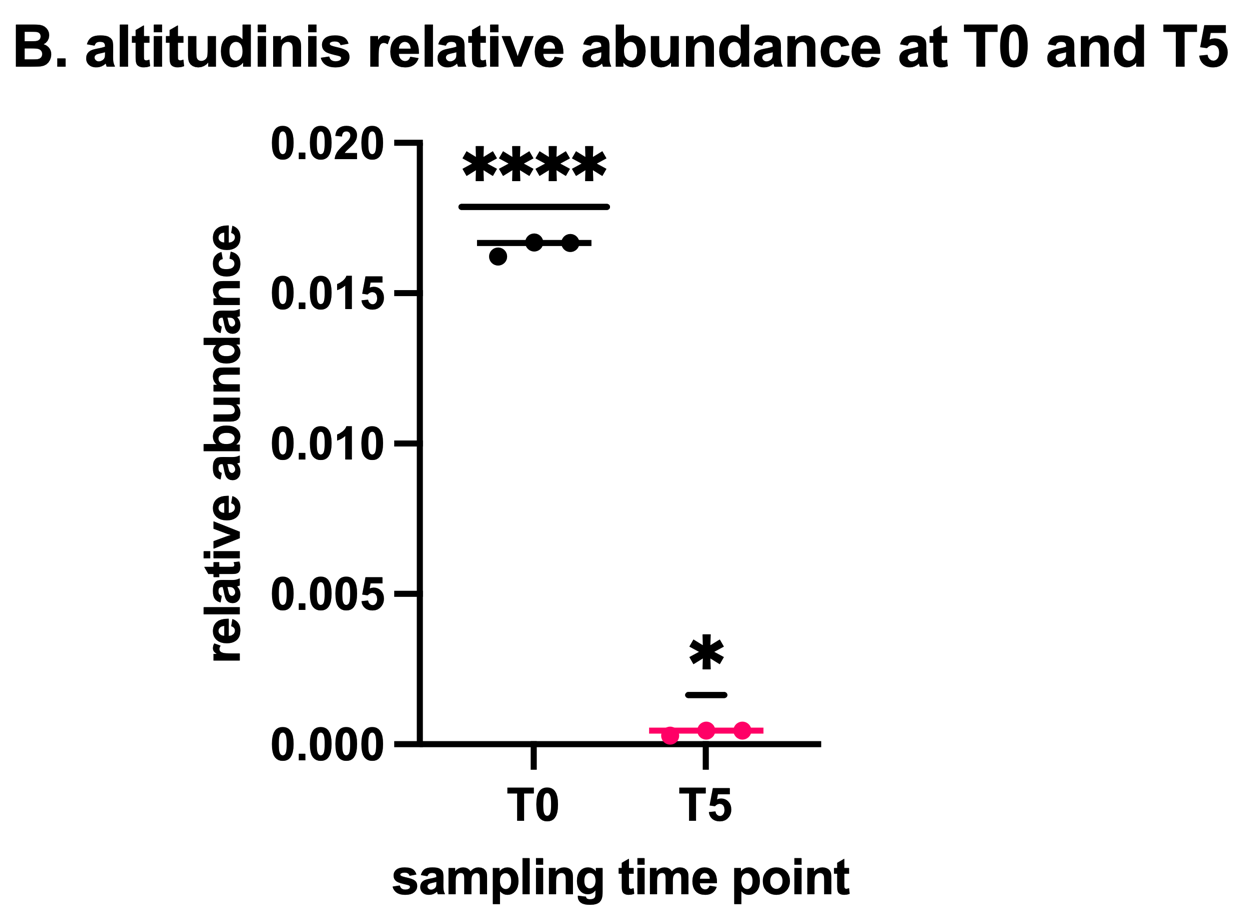

**Figure S2: Relative abundance of B. altitudinis after 0 days and 5 days incubation.** The decrease of relative abundance of B. altitudinis, when grown in the SynCom on mm9/7 agar at inoculation and after 5 days of incubation at 22 °C.

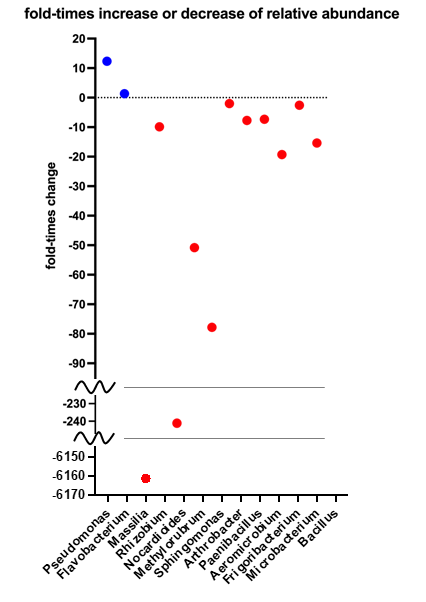

**Figure S3: Time-fold increase or decrease of relative abundance of SynCom members over incubation time** The calculation of the increase (blue) or decrease (red) was based on the relative abundance after 0 and 5 days of incubation in a whole SynCom co-culture (0.2 OD600 of each strain). The SynCom was cultured on MM9/7 agar at 22 °C and relative abundance was investigated by 16S rRNA/ITS2 MiSeq illumina amplicon sequencing.

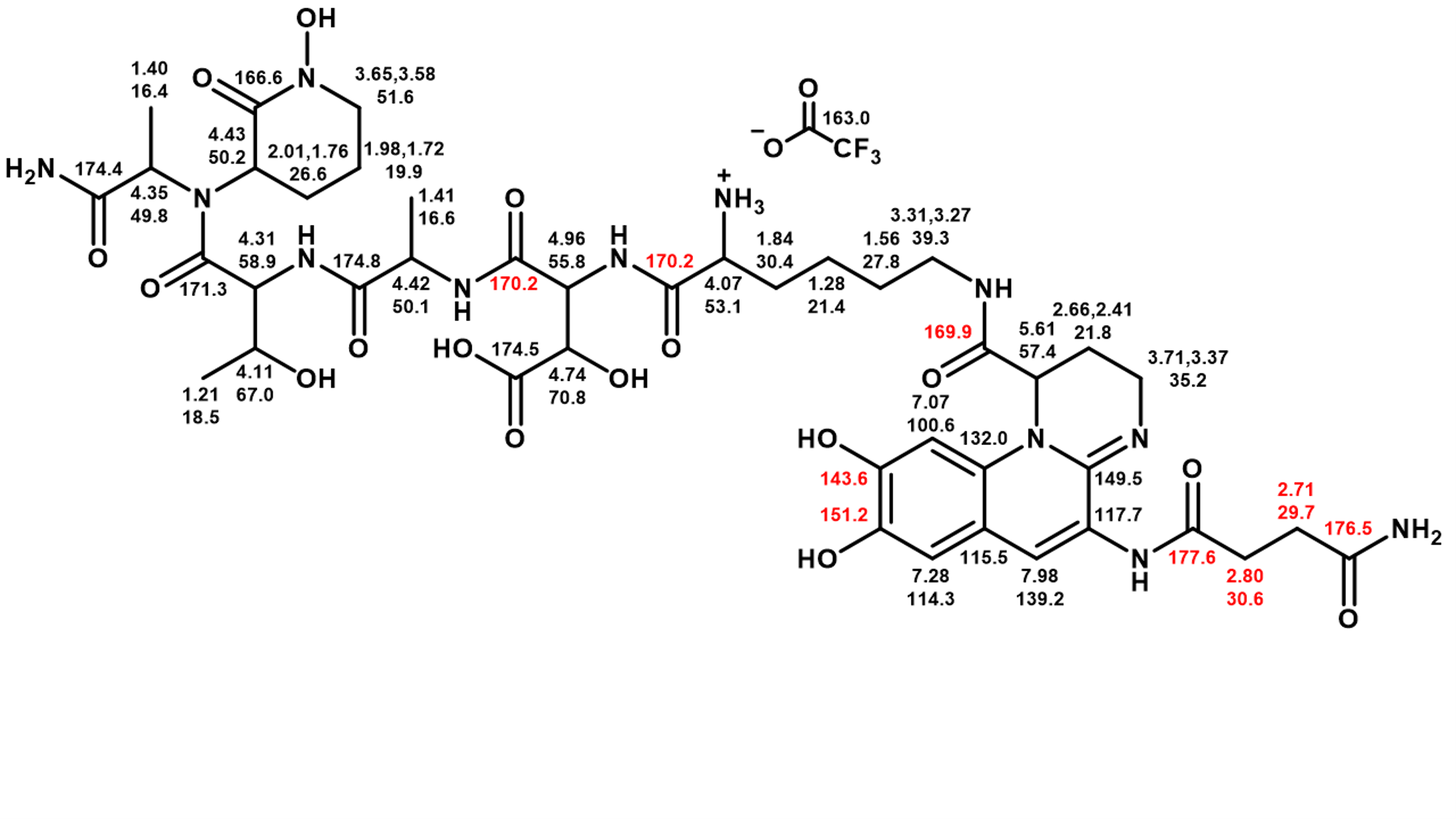

**Figure S4: Structure of pseudobactin A TFA salt (1) with 1H and 13C chemical shift assignments in D2O at 700** **MHz** NB: 1) ^1^H NMR chemical shifts are in good agreement with Teintze, M.; Leong, J. Structure of pseudobactin A, a second siderophore from plant growth promoting Pseudomonas B10. Biochemistry 1981, 20, 6457–6462. DOI:10.1021/bi00525a026. 2) Some uncertainty is associated with the assignments in red.

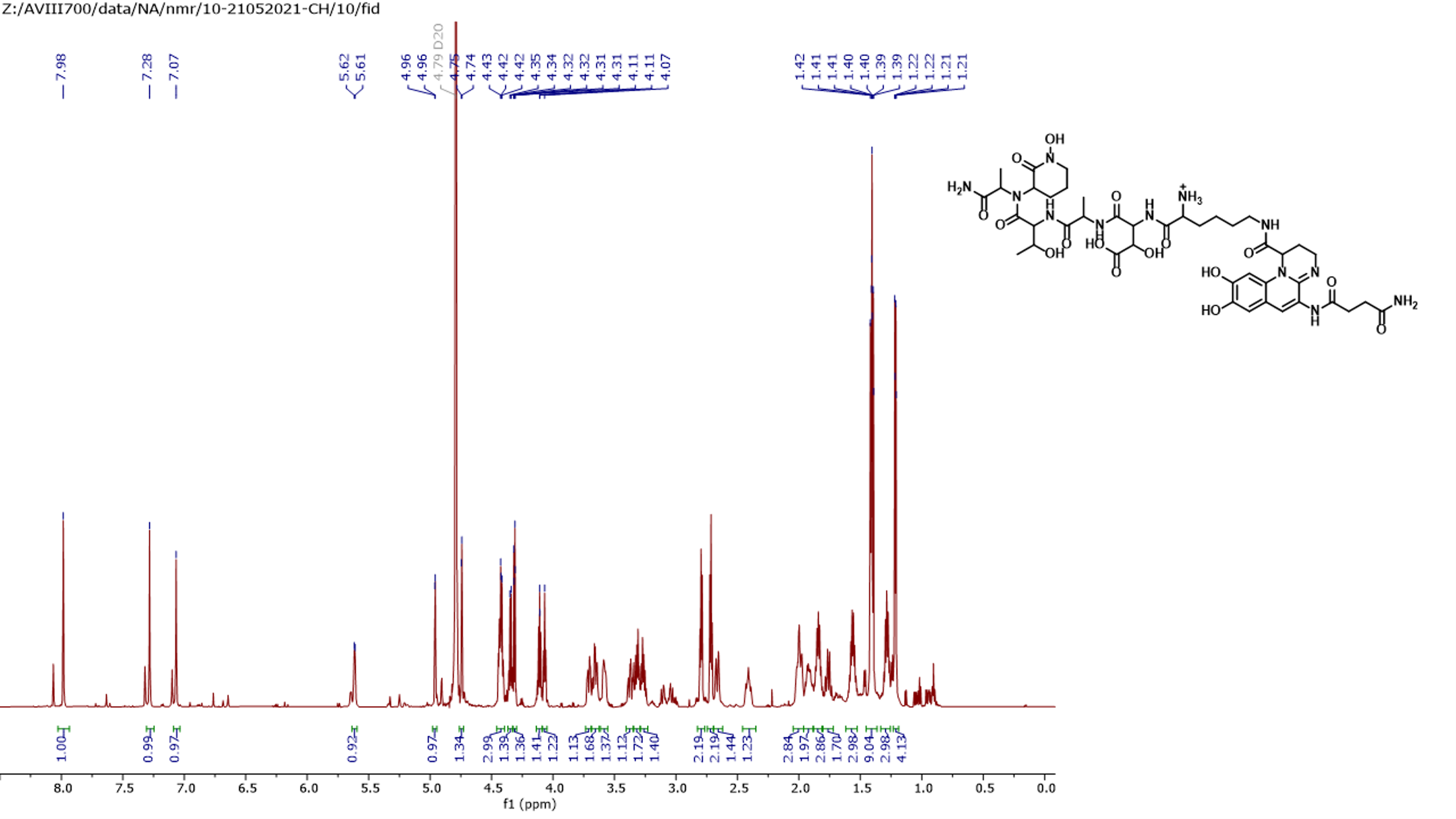

**Figure S5: ^1^H NMR (D2O, 700 MHz) of pseudobactin A TFA salt (1)**

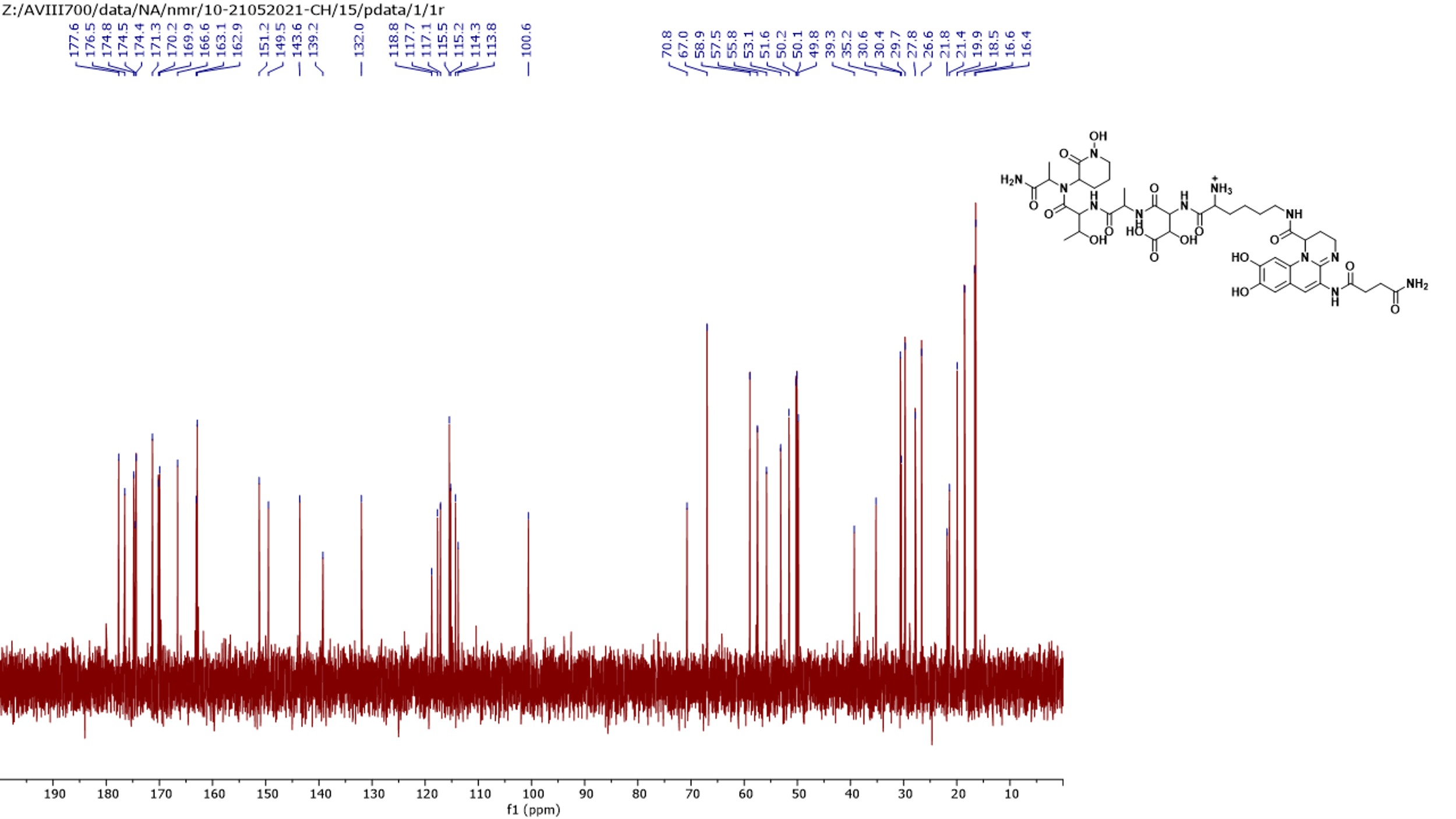

**Figure S6: ^13^C NMR (D2O, 175 MHz) of pseudobactin A TFA salt (1)**

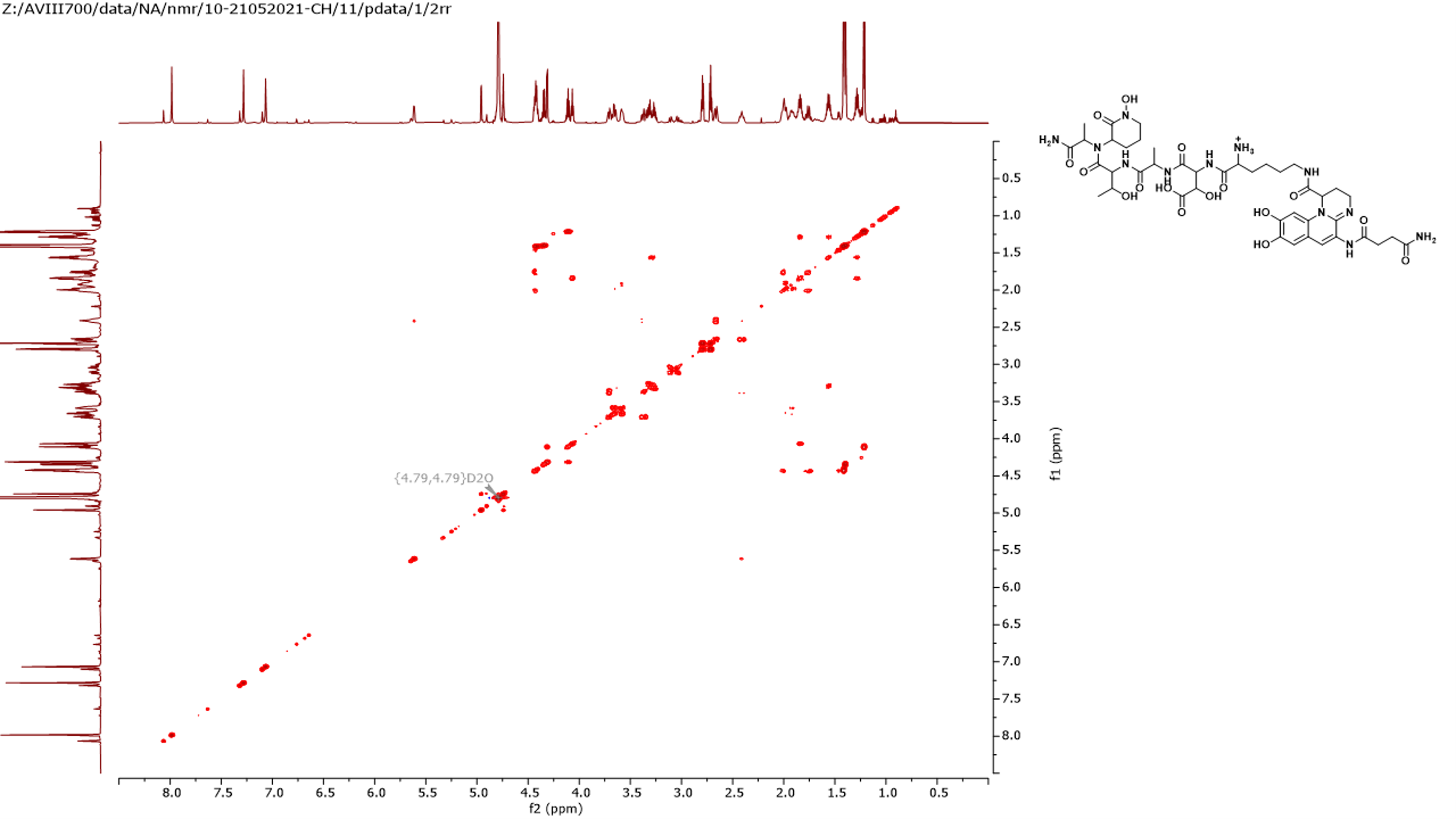

**Figure S7: COSY NMR (D2O, 700 MHz) of pseudobactin A TFA salt (1)**

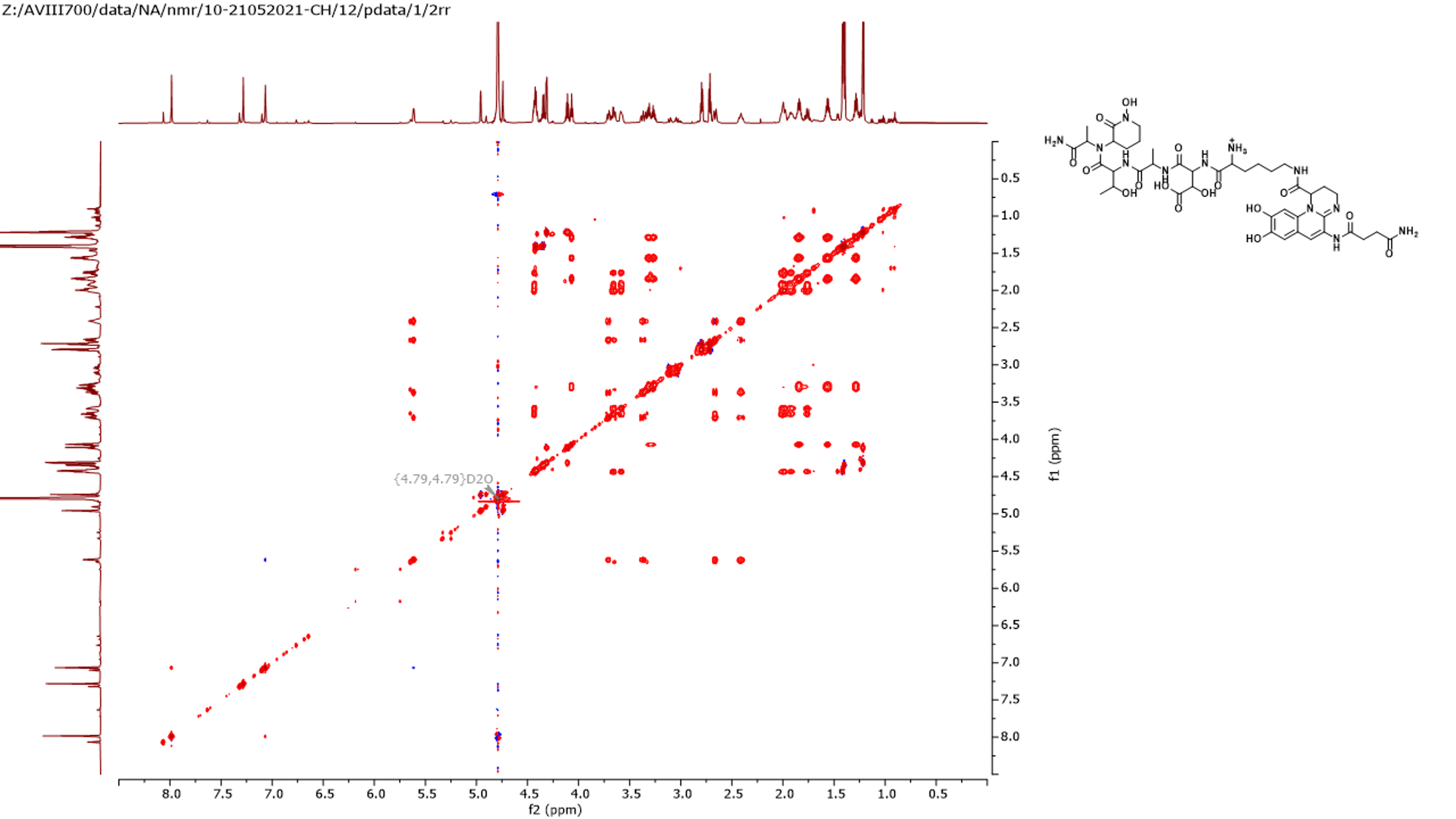

**Figure S8: TOCSY NMR (D2O, 700 MHz) of pseudobactin A TFA salt (1)**

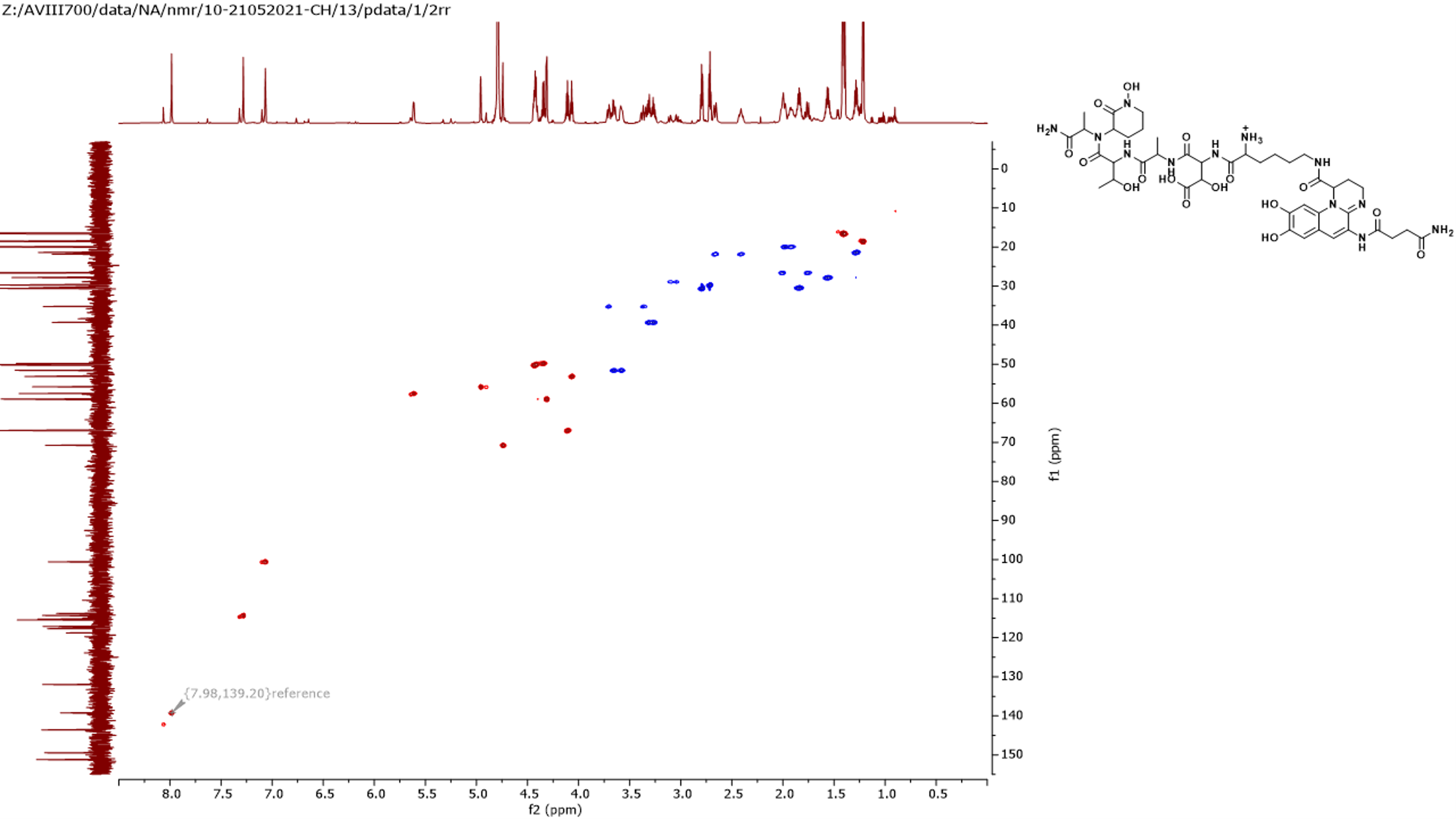

**Figure S9: Figure S5 HSQC NMR (D2O, 700 MHz) of pseudobactin A TFA salt (1)**

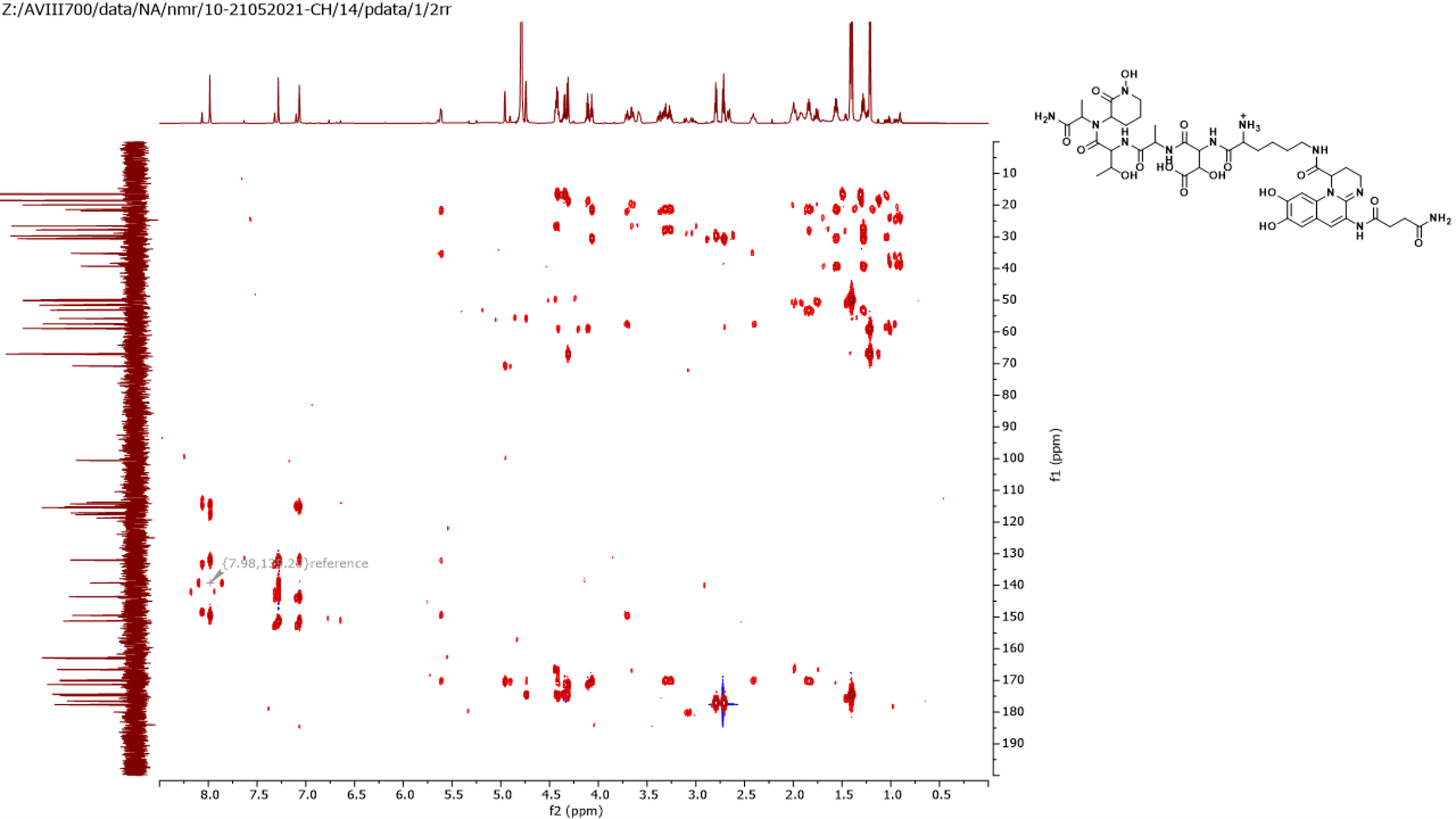

**Figure S10:HMBC NMR (D2O, 700 MHz) of pseudobactin A TFA salt (1)**

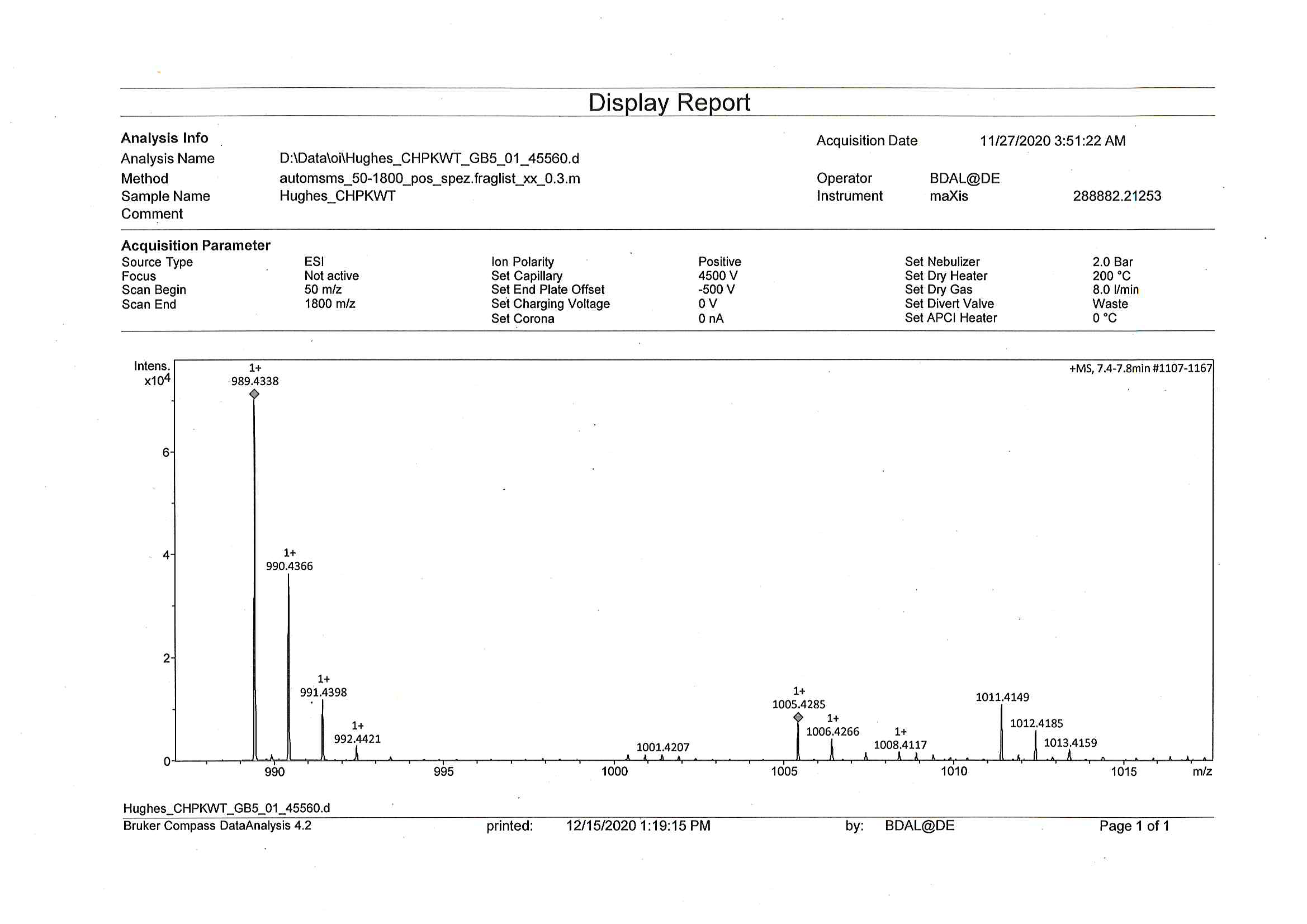

*m/z* [M+H]^+^ = 989.4338, calcd for C_42_H_61_N_12_O_16_, 989.4323

**Figure S11: HR¬MS spectrum of pseudobactin A TFA salt (1)**

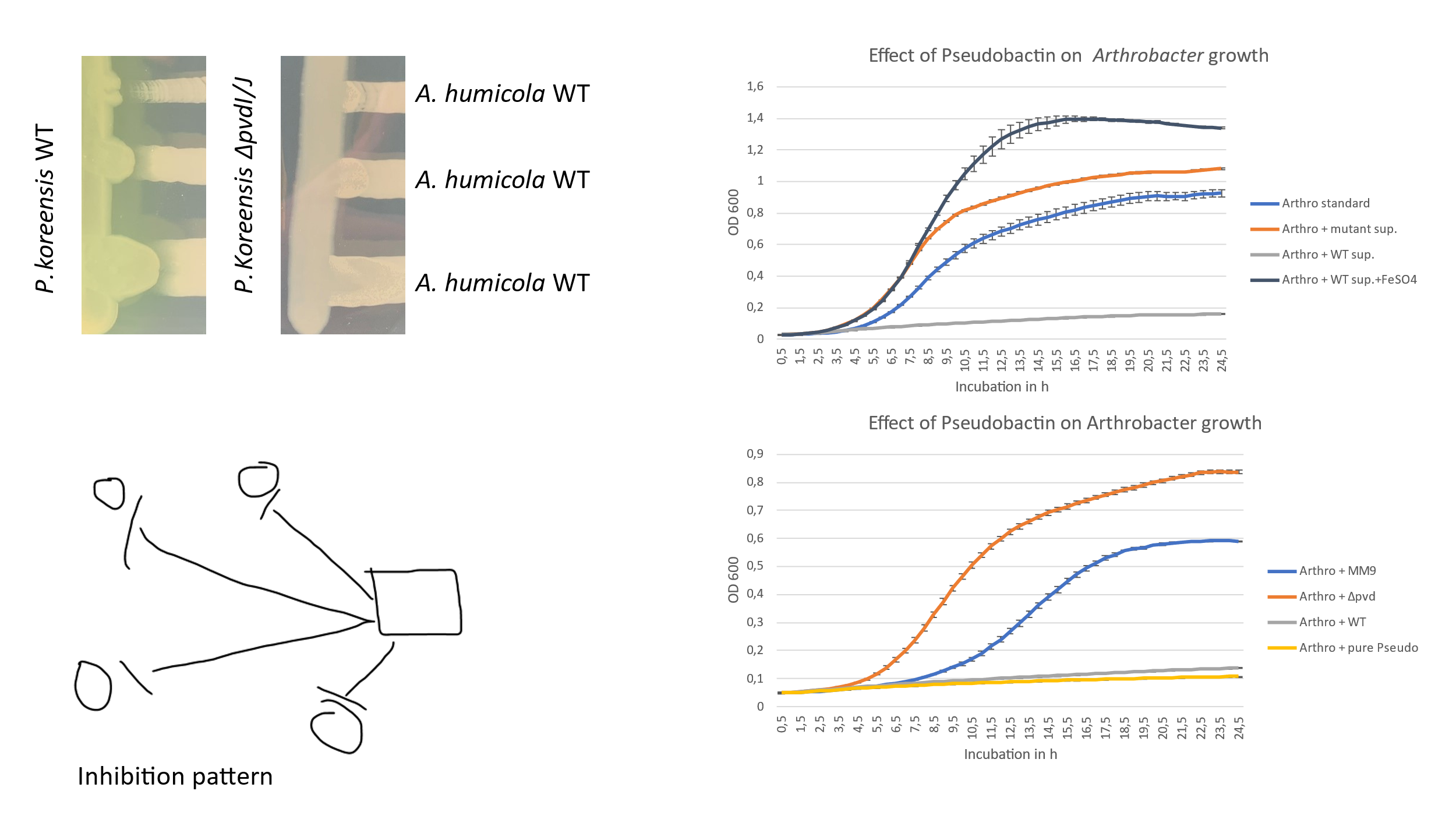

a) **Cross-streaking experiment of *P. koreensis* WT and mutant**
b) **HPLC-MS measurement of *P. koreensis* WT and mutant supernatant**

*P. koreensis* WT

*ΔpvdI/J* mutant

Pseudobactin [M+H]^+^

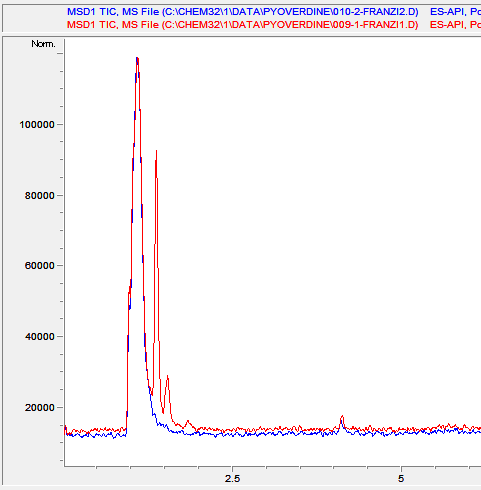

**Figure S12: Pseudobactin production and activity in P. koreensis WT and ΔpvdI/J mutant**. a) HPLC-MS of P. koreensis WT supernatant (red) and ΔpvdI/J supernatant (blue) was prepared as explained in material and methods section. A clear peak for the mass of pseudobactin is visible in WT supernatant and no peak can be seen in mutant supernatant. b) Cross-streaking experiments on f-base agar for the detection of pseudobactins´ inhibitory interaction with A. humicola. Inhibition zones and fluorescence of pseudobactin can be seen on the left (P. koreensis WT). Loss of inhibitory activity and fluorescence is observed for the mutant (right).

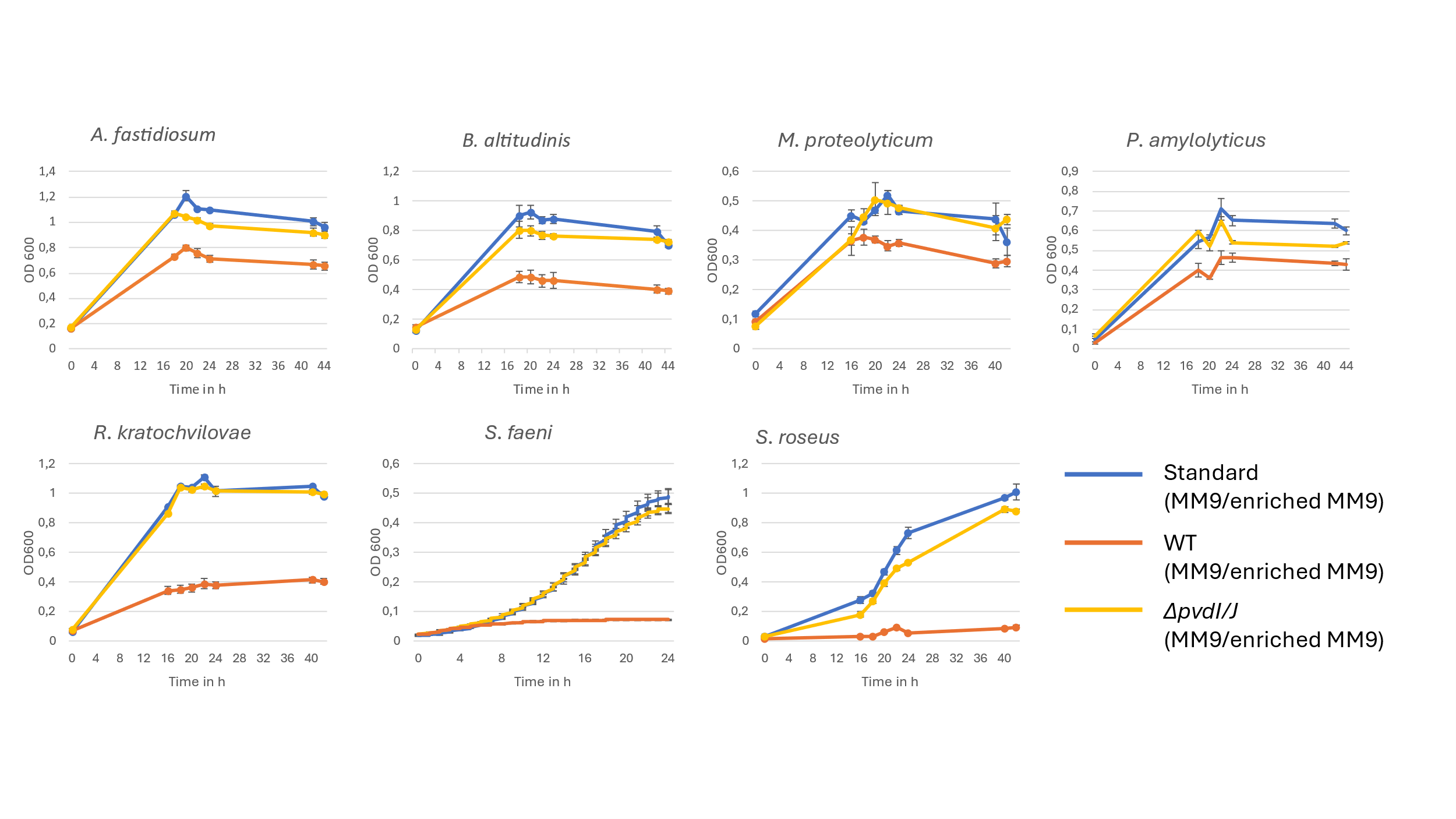

**Figure S13: Growth curves of SynCom members in presence or absence of pseudobactin** SynCom members were grown in MM9 or enriched MM9 in presence of sterile supernatant of P. koreensis WT (containing pseudobactin) and in presence of sterile supernatant of P. koreensis ΔpvdI/J mutant (no pseudobactin). Growth was observed by OD_600_ measurement at RT and 180 rpm shaking in an TECAN 2000 device. Experiments were performed in triplicates. For information on media (MM9 or enriched MM9) used for each strain see Fig. S1 and S2.

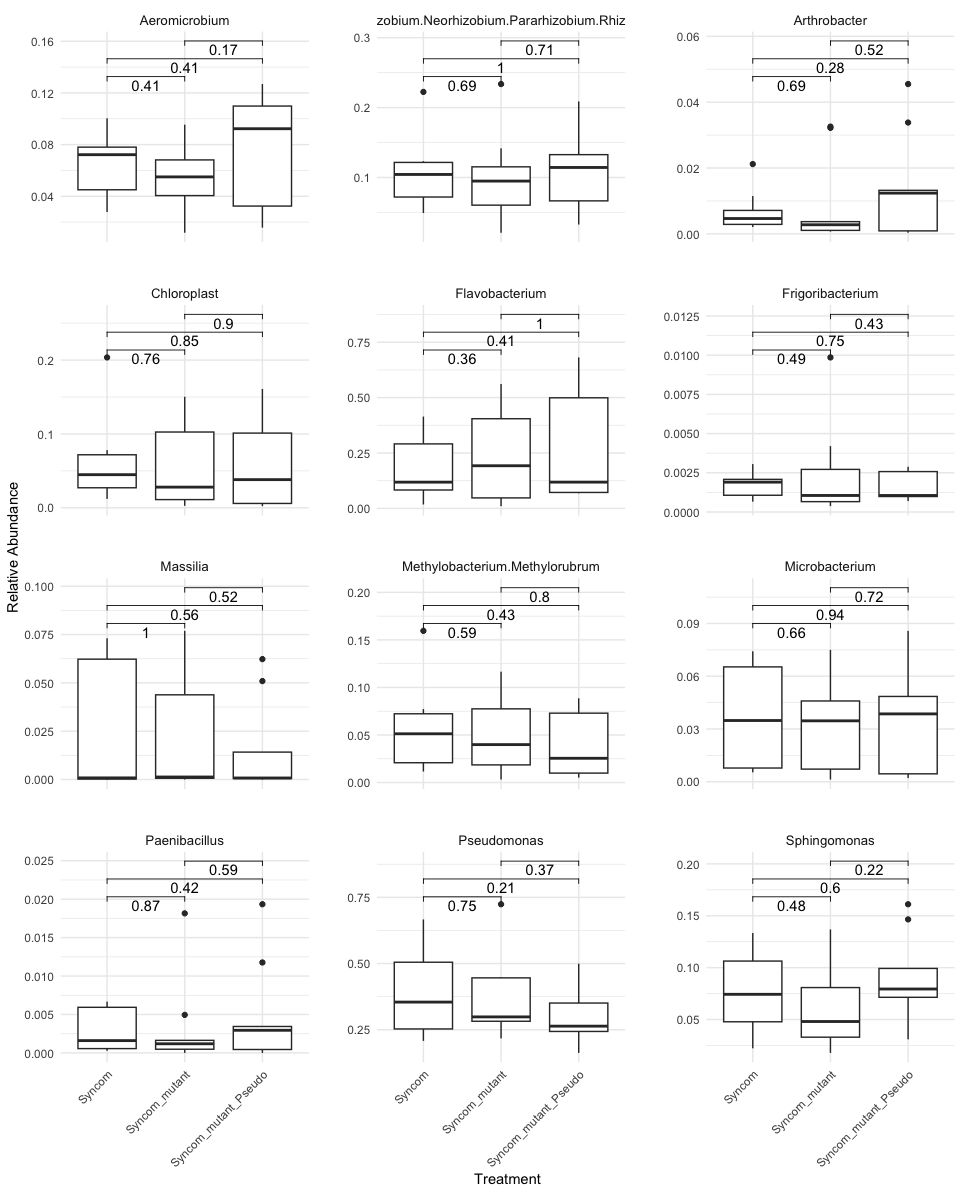

Others

**Figure S14: T-test of each SynCom bacterium for the experiment (Fig.6): The effect of pseudobactin on the SynCom composition in planta** T-test was performed to see significant changes of the relative abundance of SynCom members grown on A. thaliana with SynCom WT, SynCom mutant and SynCom pseudobactin. *No data for N. cavernae and B. altitudinis because no reads for the organisms were obtained.
